# Division of labor between ERK/MAPK and Notch signaling in regulating distinct proliferative populations during jellyfish tentacle regeneration

**DOI:** 10.64898/2026.03.08.710341

**Authors:** Sosuke Fujita, Keita Ogasawara, Ren Kanehisa, Ryo Nakajima, Hiroko Nakatani, Natsuki Shinoda, Masayuki Miura, Yu-ichiro Nakajima

## Abstract

Regeneration restores lost tissues through diverse cellular strategies that vary across species and tissue contexts. In many systems, regeneration involves blastema formation; however, how blastema formation is coordinated in tissues employing multiple stem/progenitor populations remains unclear. Here, we investigate tentacle regeneration in the hydrozoan jellyfish *Cladonema*, in which our previous work identified two distinct proliferative populations: resident homeostatic stem cells (RHSCs), which contribute to tissue production during homeostasis and regeneration, and repair-specific proliferative cells (RSPCs), which are transiently induced upon injury to form the blastema. While extensive cell death occurs shortly after injury, it is largely dispensable for blastema formation. Using pharmacological inhibition and cell proliferation analyses, we identify ERK/MAPK signaling as a key regulator of blastema formation. ERK/MAPK signaling is rapidly activated at the injury site and promotes proliferation of RSPCs. In contrast, inhibition of Notch signaling disrupts nematocyte differentiation and increases proliferative activity in RHSCs, indicating that Notch signaling governs the balance between differentiation and proliferation in RHSCs. Together, these findings reveal a division of labor between conserved signaling pathways in regulating distinct proliferative populations during tentacle regeneration.

## Introduction

Regeneration, the ability to reconstruct lost tissues or organs, varies widely across the animal kingdom. While most mammals, including humans, exhibit limited regenerative capacity, certain animals such as salamanders, planarians, and cnidarians can restore complex structures or even large portions of their bodies (Bely and Nyberg, 2010; Srivastava, 2021). Understanding the cellular and molecular mechanisms that enable such regenerative responses remains a central goal of regenerative biology. In many regenerative systems, tissue restoration is mediated not by a single homogeneous stem cell population, but by heterogeneous stem or progenitor populations with distinct developmental potentials. In planarians, for example, the neoblast compartment contains pluripotent stem cells together with molecularly and functionally distinct lineage-committed progenitors (van Wolfswinkel et al., 2014; Scimone et al., 2014; Zeng et al., 2018; Salvetti and Rossi, 2019). In contrast, the salamander limb blastema is composed of progenitor cells that retain substantial lineage restrictions during regeneration (Kragl et al., 2009; Sandoval-Guzman et al., 2014). These findings raise a broader question of how distinct stem/progenitor populations are coordinated within the same regenerating tissue and whether common or population-specific signaling mechanisms regulate their behaviors.

A number of evolutionarily conserved signaling pathways regulate regenerative proliferation, patterning, and cell-fate decisions, including Wnt/β-catenin, extracellular signal-regulated kinase (ERK)/mitogen-activated protein kinase (MAPK), TGF-β, Notch, and retinoic acid signaling (Aztekin and Storer, 2022; Min and Whited, 2023). Among these, ERK/MAPK signaling is notable for its rapid activation following tissue injury and its involvement in blastema formation and regenerative outgrowth. ERK/MAPK signaling is required during regeneration in systems including planarians and zebrafish fins (Lee et al., 2005; Tasaki et al., 2011), and recent work has shown that ERK-dependent activation of casein kinase 2 promotes blastema-cell proliferation during appendage regeneration (Zhang et al., 2024). These studies suggest that ERK/MAPK signaling represents a broadly conserved injury-responsive pathway that can be deployed to control regenerative proliferation. However, how such conserved signaling pathways regulate distinct stem/progenitor populations within the same regenerative tissue remains poorly understood.

Cnidarians provide an informative evolutionary context in which to address this question because they combine extensive regenerative capacity with stem-cell organizations that differ markedly among species and life stages (Holstein et al., 2003; Vogg et al., 2019; Macias-Munoz, 2025; Nagai and Nakajima, 2026). In polyp-stage cnidarians such as *Hydra* and *Hydractinia*, resident multipotent or pluripotent stem/progenitor populations contribute to tissue maintenance and regeneration (Gahan et al., 2016; Vogg et al., 2019). Wnt/β-catenin and ERK/MAPK signaling have been implicated in regenerative patterning and early injury responses in these systems (Hobmayer et al., 2000; Tursch et al., 2022). In hydrozoan medusae, by contrast, proliferative stem/progenitor cells are spatially restricted to defined anatomical regions. In *Clytia*, for example, interstitial stem cells and their progeny are localized in structures including the tentacle bulbs, gonads, and manubrium (Denker et al., 2008; Leclère et al., 2012). During manubrium regeneration, proliferative cells from distant tissues can also be recruited toward the regenerating organ (Sinigaglia et al., 2020). These observations illustrate the spatially organized nature of stem/progenitor populations in medusae and make them useful systems for investigating how distinct cellular populations are coordinated during regeneration.

The hydrozoan jellyfish *Cladonema pacificum* has emerged as a useful model for studying regeneration (Accorsi et al., 2024; Fujita et al., 2021). *Cladonema* tentacles regenerate rapidly after amputation, restoring both tissue architecture and differentiated cell types (Fujita et al., 2019; Fujita et al., 2023). Previous work showed that proliferative cells are concentrated in the tentacle bulb and that cell proliferation contributes to tentacle growth, morphogenesis, and regeneration (Fujita et al., 2019). More recently, we identified two spatially and functionally distinct proliferative populations that contribute to tentacle regeneration (Fujita et al., 2023). Resident homeostatic stem cells (RHSCs) are predominantly localized in the tentacle bulb and continuously contribute to tissue production during both homeostasis and regeneration, including the generation of nematocytes. In contrast, repair-specific proliferative cells (RSPCs) appear locally following injury, form the blastema, and preferentially contribute to the epithelium of the regenerated tentacle (Fujita et al., 2023). Thus, *Cladonema* provides an opportunity to investigate how a resident homeostatic proliferative population and an injury-induced proliferative population are differentially regulated within the same regenerating appendage.

How these two proliferative populations are differentially regulated during regeneration remains unknown. ERK/MAPK and Notch signaling are particularly relevant candidates because of their established roles in injury-induced proliferation and stem/progenitor cell differentiation, respectively. In cnidarians, injury rapidly activates ERK and other MAPKs in *Hydra*, where MAPK activity is required for initiation of the regenerative program (Tursch et al., 2022). Recent work further showed that sustained ERK signaling promotes activation of Wnt pathway components and supports formation of the head organizer during *Hydra vulgaris* regeneration (Juanico et al., 2026). Together with the conserved role of ERK/MAPK signaling in blastema formation in bilaterian systems, these observations raise the possibility that ERK/MAPK signaling regulates the injury-induced RSPC response in *Cladonema*. By contrast, Notch signaling has conserved roles in regulating stem/progenitor cell differentiation and cell-fate decisions. In cnidarians, perturbation of Notch signaling affects nematocyte/cnidocyte differentiation in *Hydra, Hydractinia,* and *Nematostella* (Käsbauer et al., 2007; Marlow et al., 2012; Gahan et al., 2017). Because RHSCs contribute prominently to nematocyte production in *Cladonema* tentacles (Fujita et al., 2023), these findings raise the possibility that Notch signaling regulates RHSC-associated proliferation and nematocyte differentiation in the tentacle bulb.

Here, we investigate how these conserved signaling pathways regulate the distinct proliferative populations involved in *Cladonema* tentacle regeneration. We first characterize early cellular and molecular events following tentacle amputation and then examine the roles of ERK/MAPK and Notch signaling in regenerative proliferation and differentiation. We show that ERK/MAPK signaling is rapidly activated at the injury site and preferentially promotes blastema-associated proliferation, whereas Notch signaling regulates nematocyte differentiation and proliferative activity in RHSCs. Together, our findings support a model in which tentacle regeneration is coordinated through differential regulation of spatially and functionally distinct proliferative populations.

## Materials and Methods

### Animal culture and surgical manipulation

*Cladonema pacificum* medusae (female strain 6W; Masuda-Ozawa et al., 2022; Takeda et al., 2018) were maintained in plastic containers (V-7, 85 mL; V-8, 200 mL; or V-9, 380 mL; AS ONE) filled to approximately 60–80% of their capacity with artificial seawater (ASW) at 22 °C. Under standard culture conditions, medusae were exposed to a 30-min dark period each day to induce spawning and were otherwise maintained under constant light, as previously described (Masuda-Ozawa et al., 2022; Kanehisa et al., 2026). Approximately 5–6, 10, and 15 medusae were maintained in V-7, V-8, and V-9 containers, respectively. ASW was prepared by dissolving SEA LIFE salt (Marin Tech) in tap water to a salinity of 24 ppt, followed by addition of a chlorine neutralizer (Coroline Off; GEX Co.) at 0.2 mL/L. Medusae were fed live brine shrimp (*Artemia*; A&A Marine LLC) approximately twice per week in sufficient amounts to feed all animals; the amount was not quantitatively standardized.

For surgical manipulation, medusae were anesthetized in 7% MgCl₂ prepared in deionized water for 2 min in a Petri dish, as previously described (Fujita et al., 2023). Tentacles were amputated using micro scissors under a stereomicroscope at the position indicated in Fig. 1A and 1B. Following amputation, medusae were immediately transferred back to fresh ASW and allowed to recover.

**Figure 1.**
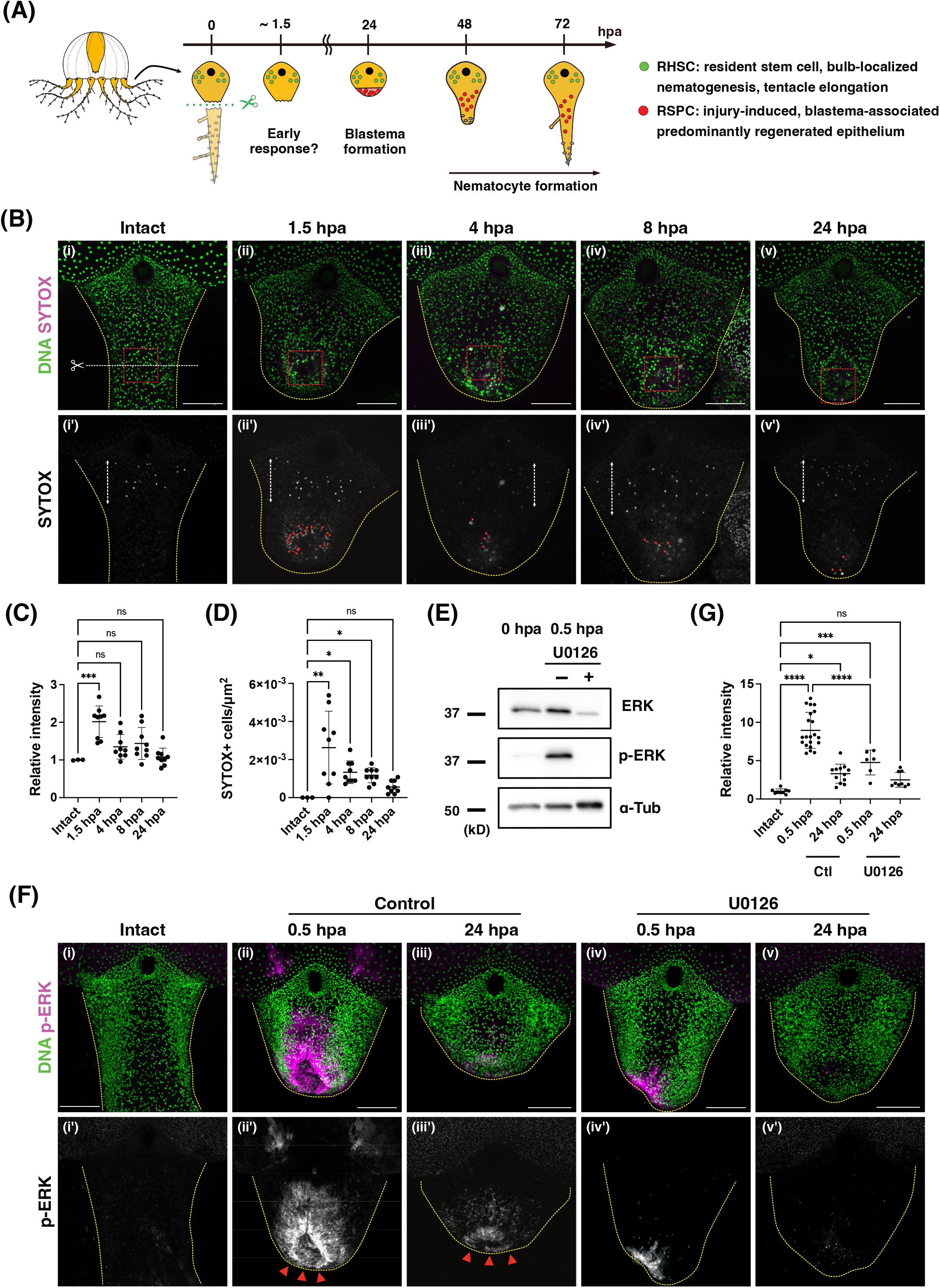
Early injury responses during tentacle regeneration include transient cell death and rapid ERK activation. **(A)** Schematic illustration of the *Cladonema* tentacle regeneration following amputation. Resident homeostatic stem cells (RHSCs; green) are predominantly localized in the tentacle bulb, whereas repair-specific proliferative cells (RSPCs; red) arise locally after injury and populate the regenerating blastema. **(B)** Detection of cell death by SYTOX Green staining. Representative images of intact and regenerating tentacles at the indicated time points. Nuclei were counterstained with Hoechst. Yellow dashed lines outline the tentacle region, red arrowheads indicate SYTOX-positive cells, and red dashed boxes indicate the fixed injury-proximal regions used for quantification in (C) and (D). White dashed arrows indicate regions of non-specific fluorescence and white dashed lines indicate the amputation position. **(C)** Quantification of relative SYTOX Green fluorescence intensity at the indicated time points, normalized to intact controls. Intact: n = 3, 1.5 hpa: n = 9; 4 hpa: n = 9; 8 hpa: n = 9; 24 hpa: n = 9. **(D)** Quantification of SYTOX Green-positive cell density within the same regions. Only SYTOX signals overlapping Hoechst-positive nuclei were counted, and obvious non-specific fluorescence was excluded from the analysis. Sample numbers are identical to those shown in (C). **(E)** Western blot analysis of phosphorylated ERK (p-ERK) and total ERK in tentacle lysates collected at the indicated time points. Where indicated, medusae were treated with the MEK inhibitor U0126. α-Tubulin was used as a loading control. **(F)** Immunofluorescence staining for p-ERK in intact and regenerating tentacles at 0.5 and 24 hpa. p-ERK signal was rapidly induced at the injury site following amputation and remained detectable at 24 hpa. Medusae were treated with 10 μM U0126 immediately after amputation until the indicated time point. Representative maximum-intensity projections of z-stacks are shown. Red arrowheads indicate regions enriched for p-ERK signals. **(G)** Quantification of relative p-ERK fluorescence intensity at the injury site based on images shown in (F). Injury-induced p-ERK intensity at 0.5 hpa was significantly reduced by U0126 treatment. Intact: n = 9; control 0.5 hpa: n = 21; control 24 hpa: n = 12; U0126 0.5 hpa: n = 6; U0126 24 hpa: n = 9. Data are shown as mean ± SD. Statistical significance was assessed by one-way ANOVA followed by Dunnett’s multiple-comparison test in (C), Kruskal–Wallis test followed by Dunn’s multiple-comparison test in (D), and one-way ANOVA followed by Tukey’s multiple-comparison test in (G). Scale bars: (B), (F) 100 μm.

### Drug treatments

Medusae were incubated in ASW containing the following reagents: 10 mM hydroxyurea (HU; 085–06653, Wako), 5, 10, or 25 μM Q-VD-Oph (S7311; Selleck Chemicals), 1, 10, or 20 μM Z-DEVD-FMK (S7312; Selleck Chemicals), 0.1 μM or 1 μM PD184352 (165-26761; FUJIFILM Wako Pure Chemical Corporation), 1 μM or 10 μM U0126 (662005; Merck Millipore), 10 μM DAPT (043-33581; FUJIFILM Wako Pure Chemical Corporation), or 5 μM LY411575 (L0359; Tokyo Chemical Industry). All compounds were administered individually and were not used in combination. HU was dissolved directly in ASW, whereas Q-VD-Oph, Z-DEVD-FMK, PD184352, U0126, DAPT, and LY411575 were prepared as stock solutions in DMSO. ASW alone was used as the control for HU treatment, whereas control animals for the other drug treatments received the corresponding concentration of DMSO. For each inhibitor, concentrations were selected based on preliminary titration experiments to identify doses that produced the intended biological effect without overt morphological abnormalities or lethality under the conditions used for each assay.

For caspase-inhibitor experiments, medusae were pretreated with Q-VD-Oph or Z-DEVD-FMK for 1.5 h before tentacle amputation. Inhibitor treatment was continued until 1.5 hpa, resulting in a total treatment period of 3 h. After 1.5 hpa, medusae were transferred to fresh ASW without inhibitor and allowed to recover until the indicated experimental endpoint. For analysis of early cell death, 10 μM Q-VD-Oph and 1 μM Z-DEVD-FMK were used. For the subsequent proliferation assay, higher concentrations (25 μM Q-VD-Oph and 20 μM Z-DEVD-FMK) were used during the same transient treatment window to provide a more stringent pharmacological test of whether early caspase inhibition affects subsequent proliferation. Cell proliferation was analyzed at 24 hpa. For regenerative-outgrowth assays, Q-VD-Oph was tested at 5 and 10 μM and Z-DEVD-FMK at 1 and 10 μM, with regenerative tentacle length measured at later time points up to 96 hpa.

For MEK- and γ-secretase-inhibitor experiments, treatment was initiated immediately after tentacle amputation and continued until the indicated experimental endpoint. Regenerating medusae were treated until 24 hpa for proliferation analyses and throughout the indicated multi-day period for analyses of nematocyte formation and regenerative outgrowth. Intact medusae were treated only in experiments specifically examining proliferation under homeostatic conditions. During multi-day treatments, drug-containing ASW was renewed every 48 h.

### SYTOX staining

Intact or tentacle-amputated medusae were incubated for 30 min in ASW containing 2 μM SYTOX Green Nucleic Acid Stain (Thermo Fisher Scientific, S7020) and Hoechst 33342. After incubation, samples were washed three times with ASW and mounted on slides in 7% MgCl₂ for imaging. For quantification of SYTOX-positive cells, only SYTOX signals overlapping Hoechst-positive nuclei were counted. Quantification was performed within a fixed ROI positioned around the injury-proximal/blastema region, thereby excluding obvious non-specific fluorescence from the eye spot and tentacle bulb.

### TUNEL assay

TUNEL staining was performed using the Click-iT™ Plus TUNEL Assay Kit for In Situ Apoptosis Detection (Thermo Fisher Scientific, C10618), according to the manufacturer’s instructions. Briefly, *Cladonema* medusae were anesthetized in 7% MgCl₂ prepared in deionized water for 5 min and fixed in 4% paraformaldehyde (PFA) in ASW for 1 h at room temperature or overnight at 4 °C. Samples were then washed and subjected to TUNEL labeling to detect apoptotic cells.

### Immunofluorescence staining

Medusae were anesthetized in 7% MgCl₂ for 5 min and fixed in 4% PFA in ASW for 1 h at room temperature or overnight at 4 °C. Samples were washed with phosphate-buffered saline (PBS) containing 0.1% Triton X-100 (0.1% PBT). No separate blocking step was performed. Samples were incubated overnight at 4 °C with primary antibodies diluted in 0.1% PBT.

After washing, samples were incubated for 1 h at room temperature in the dark with secondary antibodies (1:500; Alexa Fluor 488, 555, or 647 donkey or goat anti-mouse or anti-rabbit IgG; Thermo Fisher Scientific, A21245, A21236, A31570, A31572, A21202, A21206) and Hoechst 33342 (1:500; Thermo Fisher Scientific, H1399). Samples were washed three times in 0.1% PBT and mounted in 70% glycerol.

The following primary antibodies were used: rabbit anti–phospho-histone H3 (Ser10) (1:500; Upstate, 06–570), mouse anti–α-tubulin (1:500; Sigma-Aldrich, T6199), and rabbit anti–phospho-p44/42 MAPK (ERK1/2) (1:100; Cell Signaling Technology, #4370).

For pERK immunostaining only, fluorescence signals were amplified using a tyramide signal amplification (TSA) protocol adapted from Kanehisa et al. (2026). Following primary antibody incubation and washing, samples were incubated overnight with Rabbit MAX PO (1:20; Nichirei Corporation), corresponding to the host species of the primary antibody, in 0.5% blocking buffer (0.5% blocking reagent [Roche] in Maleic acid buffer). Samples were then washed three times with 0.1% PBT and three times with TNT buffer (100 mM NaCl, 100 mM Tris–HCl, pH 9.5, 0.1% Tween-20), followed by incubation with Cy3- or Cy5-tyramide solution (TSA Plus Cyanine 3 or 5; Akoya Biosciences) for 10 min.

For visualization of mature nematocytes, nuclear DNA and poly-γ-glutamate were stained with DAPI (1:250; Invitrogen, D1306) instead of Hoechst 33342 (Szczepanek et al., 2002).

### Protein extraction and western blotting

For western blot analysis, four tentacles from each of five medusae were pooled per sample and lysed in 50 μL of lysis buffer supplemented with 1× cOmplete ULTRA EDTA-free protease inhibitor cocktail (Cat#05892953001, Roche). Samples were homogenized and centrifuged at 20,380 × g for 15 min at 4 °C, and the supernatants were collected.

Protein concentrations were determined using the Pierce BCA Protein Assay Kit (Thermo Fisher Scientific, 23225). Samples were mixed with 6× Laemmli sample buffer to a final concentration of 1× and boiled at 98 °C for 5 min. Proteins were separated by SDS-PAGE and transferred onto PVDF membranes (Immobilon-P, IPVH00010; Millipore). Membranes were blocked in 5% skim milk in TBST (Tris-buffered saline containing 0.1% Tween-20) and incubated with primary antibodies diluted in blocking solution.

The following primary antibodies were used: rabbit anti-p44/42 MAPK (ERK1/2) (1:1000; Cell Signaling Technology, #9102), rabbit anti-phospho-p44/42 MAPK (ERK1/2) (1:1000; Cell Signaling Technology, #4370), and mouse anti-α-tubulin (DM1A; 1:5000; Sigma-Aldrich, T9026) as a loading control.

Secondary antibodies were HRP-conjugated goat anti-rabbit IgG (1:10,000; Cell Signaling Technology, 7074S) and HRP-conjugated anti-mouse IgG (1:10,000; Promega, W402B). Signals were detected using Immobilon Western Chemiluminescent HRP Substrate (WBKLS0500; Millipore) and imaged with a FUSION SOLO.7S.EDGE system (Vilber-Lourmat).

### EdU labeling

To label S-phase cells, medusae were treated as described previously (Fujita et al., 2022; Fujita et al., 2019). Briefly, medusae were incubated in ASW containing 150 μM 5-ethynyl-2′-deoxyuridine (EdU; Invitrogen, C10340) for 1 h. After incubation, medusae were anesthetized with 7% MgCl₂ and fixed in 4% PFA in ASW for 1 h at room temperature or overnight at 4 °C. Samples were washed with 0.1% PBT and incubated with the EdU reaction cocktail (1× reaction buffer, CuSO₄, Alexa Fluor azide 647, and reaction buffer additive; Invitrogen, C10340) for 30 min in the dark.

After the EdU reaction, samples were washed and counterstained with Hoechst 33342 in 0.1% PBT for 30 min. When combined with immunofluorescence, the EdU reaction was performed after secondary antibody incubation.

### Fluorescence image acquisition & Image analysis

Confocal images were acquired using a Zeiss LSM 880 confocal microscope. For all fluorescence imaging, z-stacks were acquired and displayed as maximum-intensity projections. Image processing and quantification were performed using ImageJ/Fiji.

For cell counting and fluorescence-intensity measurements, a fixed square region of interest (ROI; 74.72 × 74.72 µm) was defined using the Rectangle tool and positioned within the indicated region of the tentacle. The same ROI size was used consistently across samples. For quantification of EdU-, PH3-, and SYTOX-positive cells, positive cells within each ROI were manually counted. Cell density was calculated as the number of positive cells divided by the ROI area and expressed as cells/µm². For fluorescence-intensity measurements, an ROI of the same size was used, and the mean gray value within each ROI was measured using the Measure function in Fiji.

### Statistical analysis

All experiments were performed with at least two independent biological replicates. The number of animals used in each biological replicate varied among experiments and is specified in the corresponding figure legends. Statistical analyses were performed using GraphPad Prism9. Microsoft Excel was used for data organization. Statistical comparisons were performed using two-tailed unpaired t-tests, one-way ANOVA followed by Dunnett’s, Tukey’s, or Šídák’s multiple-comparison tests, Kruskal–Wallis tests followed by Dunn’s multiple-comparison test, ordinary two-way ANOVA followed by Dunnett’s multiple-comparison test, or two-way repeated-measures ANOVA with Geisser– Greenhouse correction followed by Dunnett’s multiple-comparison test, as appropriate. The statistical test used for each experiment is specified in the corresponding figure legends. Significance was indicated in the figures as follows: *p < 0.05, **p < 0.01, ***p < 0.001, ****p < 0.0001, not significant (n.s.): p ≥ 0.05. Data are presented as mean ± standard deviation (SD).

## Results

### Early events in tentacle regeneration: cell death and ERK/MAPK activation

After amputation, *Cladonema* tentacles first undergo wound healing, followed by the formation of blastema that contains undifferentiated cells within 24 hours (Fujita et al., 2023; Fig. 1A). To investigate the cellular responses and regulatory factors involved in blastema formation, we examined early regenerative events prior to the blastema appearance. During regeneration of various animal organs and tissues, cell death has been observed as an early response after injury, in addition to wound repair (Adell et al., 2025; Guerin et al., 2021; Vriz et al., 2014). In the cnidarian polyp-type animal *Hydra*, for instance, both apoptosis and caspase-dependent pyroptotic cell death occur following injury (Chen et al., 2023a; Chera et al., 2009; Chera et al., 2011; Tursch et al., 2022). Genes predicted to encode caspase-3 and gasdermin (GSDM) are also present in *Cladonema*, and the predicted proteins retain conserved features associated with caspase activity and gasdermin cleavage (Chen et al., 2023b; Sup. Figs. S1, S2). SYTOX staining revealed a significant increase in SYTOX-positive cells around the injury site at 1.5 hours post-amputation (hpa) (Fig. 1B–1D). TUNEL staining independently detected an increase in apoptotic cells at 1.5 hpa (Sup. Fig. S3A, S3B). Subsequently, the SYTOX-positive cell-death response gradually subsided and was largely resolved by 24 hpa (Fig. 1B-1D), coinciding with the timing of blastema formation (Fig. 1A). These observations indicate that cell death is rapidly induced near the amputation site during the early phase of tentacle regeneration.

Activation of the ERK/MAPK pathway is another evolutionarily conserved response to tissue injury observed across a wide range of species from bilaterians to cnidarians (DuBuc et al., 2014; Tasaki et al., 2011; Tursch et al., 2022; Wen et al., 2022). In *Hydra*, injury-induced reactive oxygen species (ROS) and Ca^2+^ signaling contribute to ERK/MAPK activation, leading to rapid phosphorylation of ERK (p-ERK) after injury (Tursch et al., 2022). Using an anti-phospho-ERK antibody, we confirmed that ERK activation occurs as early as 0.5 hpa in *Cladonema* tentacles by western blotting (Fig. 1E). Immunostaining further revealed strong p-ERK signals around the injury site (Fig. 1F). While p-ERK levels were highest shortly after amputation (Fig. 1Fi, 1Fii, 1G), p-ERK remained detectable for at least 24 hpa, albeit at reduced levels (Fig. 1Fiii). The predicted *Cladonema* ERK protein harbors a conserved TEY phosphorylation motif (Sup. Fig. S4). Pharmacological inhibition of MEK using U0126 led to a significant reduction in injury-induced ERK activation (Fig. 1F, 1G). Similarly, the second MEK inhibitor PD184352 significantly reduced p-ERK intensity at 0.5 hpa, although residual p-ERK remained detectable (Sup. Fig. S5). Together, these results show that cell death and ERK/MAPK activation are rapidly induced during the initial stages of tentacle regeneration.

In *Hydra* head regeneration, injury-induced MAPK signaling has been linked to early cell-death and Wnt responses (Chera et al., 2011; Tursch et al., 2022), while apoptotic cells can promote proliferation of neighboring cells through Wnt signaling (Chera et al., 2009). Because both ERK/MAPK activation and cell death were rapidly induced after tentacle amputation in *Cladonema*, we asked whether these two early injury responses are functionally connected and contribute to subsequent blastema proliferation.

To test whether ERK/MAPK activation affects the early cell-death response, we examined cell death at 1.5 hpa following treatment with the MEK inhibitors U0126 and PD184352. Neither inhibitor significantly altered SYTOX-positive cell death compared with controls, indicating that MEK inhibition did not detectably affect the early injury-induced cell-death response under these conditions (Sup. Fig. S6). We next asked whether this early cell-death response contributes to subsequent blastema proliferation.

### Caspase-mediated cell death does not induce compensatory proliferation during tentacle regeneration

In some regenerative systems, cell death has been proposed to promote regeneration by inducing compensatory proliferation in surrounding cells or by contributing to tissue remodeling (Adell et al., 2025; Bergmann, 2025; Guerin et al., 2021; Kawamoto et al., 2016; Vriz et al., 2014). Secreted signals released from dying cells stimulate proliferation in neighboring cells during zebrafish fin regeneration and *Hydra* head regeneration, ultimately contributing to regenerative proliferation and blastema formation (Chera et al., 2009; Gauron et al., 2013). To assess whether caspase-mediated cell death influences cell proliferation during *Cladonema* tentacle regeneration, we inhibited caspase activity using the pan-caspase inhibitor Q-VD-Oph as well as the caspase-3– targeting inhibitor Z-DEVD-FMK. Both Q-VD-Oph and Z-DEVD-FMK significantly reduced early cell death (Fig. 2A-2C) and were therefore used in subsequent experiments. To selectively perturb the early cell-death response, inhibitors were applied from 1.5 h before amputation until 1.5 hpa and were then removed.

**Figure 2.**
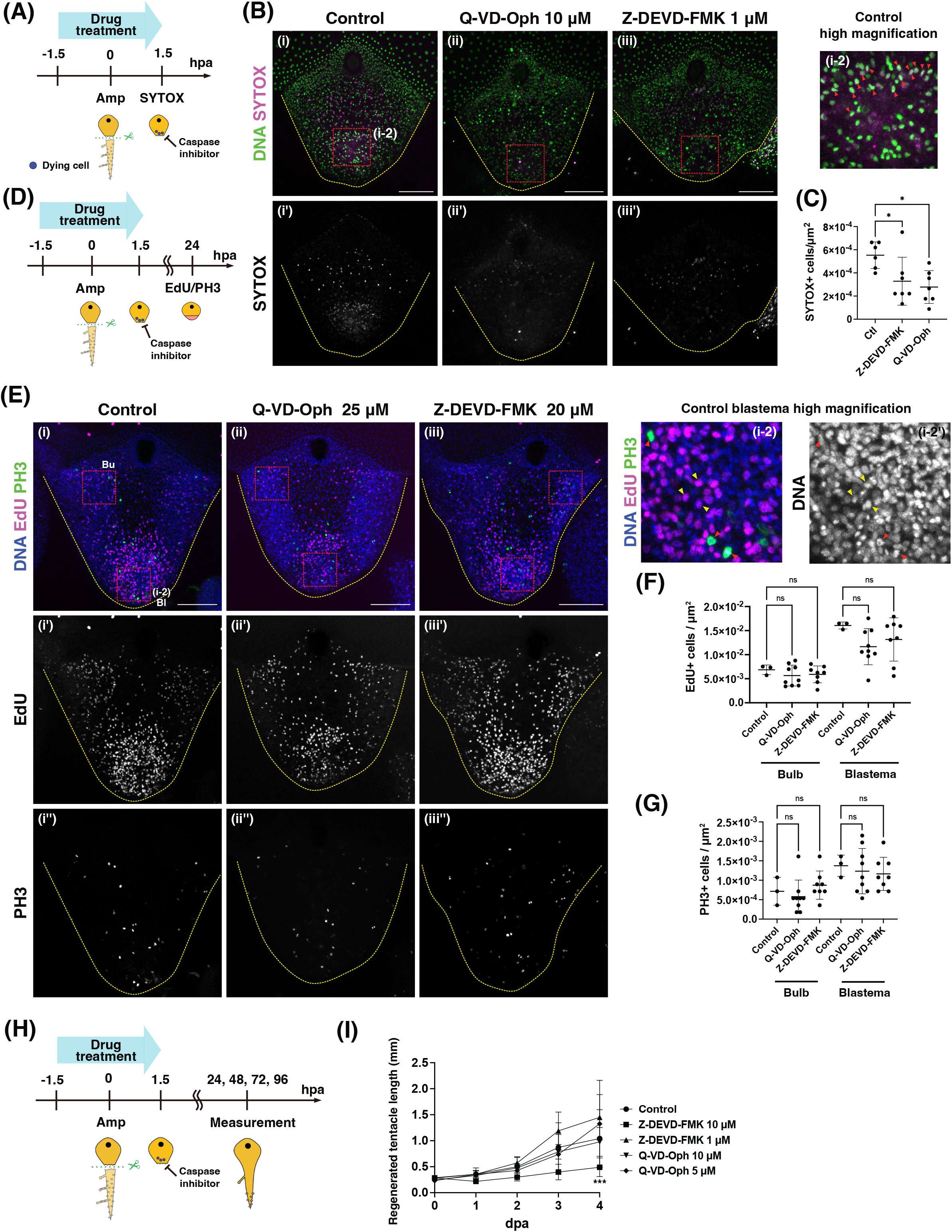
Caspase-mediated cell death is dispensable for blastema proliferation. **(A)** Experimental scheme for pharmacological inhibition of caspase activity during the early stage of tentacle regeneration. Caspase inhibitors were applied from 1.5 h before amputation until 1.5 hpa and were then removed. **(B)** SYTOX Green staining showing cell death in control and caspase inhibitor-treated tentacles at 1.5 hpa. Animals were treated with 10 μM Q-VD-Oph or 1 μM Z-DEVD-FMK. Representative maximum-intensity projections of z-stacks are shown. Nuclei were counterstained with Hoechst. Red dashed boxes indicate the wound-proximal region. Red arrowheads in the high-magnification image indicate SYTOX-positive cells overlapping with Hoechst-positive nuclei. **(C)** Quantification of SYTOX Green-positive cell density in control and caspase inhibitor-treated tentacles at 1.5 hpa. Control: n = 6; Q-VD-Oph 10 μM: n = 7; Z-DEVD-FMK 1μM: n = 7. **(D)** Experimental scheme for assessing proliferative activity during tentacle regeneration. Following transient caspase inhibitor treatment from 1.5 h before amputation until 1.5 hpa, proliferative activity was examined by EdU pulse labeling and anti-PH3 staining at 24 hpa. **(E)** Detection of S-phase cells by EdU pulse labeling and M-phase cells by PH3 staining in regenerating tentacles. Animals were treated with 25 μM Q-VD-Oph or 20 μM Z-DEVD-FMK. Representative maximum-intensity projections of z-stacks are shown. Red dashed boxes indicate the regions used for quantification; Bl, blastema; Bu, tentacle bulb. Yellow arrowheads indicate EdU-positive cells, and red arrowheads indicate PH3-positive cells in the high-magnification images, showing overlap with nuclear staining. **(F, G)** Quantification of EdU-positive (F) and PH3-positive (G) cell density in the bulb or blastema region. Control: n = 3; Q-VD-Oph, 25 μM: n = 8; Z-DEVD-FMK, 20 μM: n = 9. **(H)** Experimental scheme for assessing regenerative growth following transient caspase inhibitor treatment. Caspase inhibitors were applied from 1.5 h before amputation until 1.5 hpa and were then removed. Regenerated tentacle length was measured at 24, 48, 72, and 96 hpa. **(I)** Quantification of regenerating tentacle length following transient caspase inhibitor treatment. Control: n = 11, Z-DEVD-FMK, 10 μM: n = 8; Z-DEVD-FMK, 1 μM: n = 8, Q-VD-Oph, 10 μM: n = 8; Q-VD-Oph 5 μM: n = 8. A modest but statistically significant reduction in tentacle length was observed in animals treated with 10 μM Z-DEVD-FMK compared with controls. Representative fluorescence images in (B) and (E) are maximum-intensity projections of z-stacks. Data are shown as mean ± SD. Statistical significance was assessed by one-way ANOVA followed by Dunnett’s multiple-comparison test in (C), by one-way ANOVA followed by Tukey’s multiple-comparison test in (F) and (G), and by two-way ANOVA followed by Dunnett’s multiple-comparison test in (I). Scale bars: (B), (E) 100 μm.

To investigate the impact of caspase inhibition on regenerative proliferation, we examined cell proliferation at the injury site using EdU pulse labeling for S-phase cells and immunostaining with an anti-phospho-histone H3 (PH3) antibody to detect mitotic cells. Because EdU labels cells that enter or remain in S-phase during the 1-h pulse, whereas PH3 marks the comparatively shorter M phase, the number of EdU-positive cells was higher than that of PH3-positive cells under these conditions. For this assay, we used higher inhibitor concentrations (25 µM Q-VD-Oph and 20 µM Z-DEVD-FMK) during the same transient treatment window to provide a more stringent pharmacological test of whether early caspase inhibition affects subsequent proliferation. The densities of EdU-positive and PH3-positive cells within the blastema region of inhibitor-treated animals were comparable to those in controls (Fig. 2D-2G), indicating that transient inhibition of the early caspase-mediated cell-death response did not detectably reduce blastema proliferation at 24 hpa. In addition, transient caspase inhibitor treatment had no detectable effect on proliferative activity in the RHSC-enriched tentacle bulb (Fig. 2E-2G). Thus, even under these higher-dose pharmacological conditions, neither S-phase nor mitotic activity was detectably altered.

Given that inhibition of cell death in *Hydra* leads to incomplete head regeneration (Chen et al., 2023a; Chera et al., 2009), we further tested whether transient inhibition of early caspase-mediated cell death affects subsequent regenerative outgrowth by measuring tentacle elongation. At the concentrations shown to significantly suppress early cell death (10 µM Q-VD-Oph and 1 µM Z-DEVD-FMK), tentacle elongation was not significantly affected. Consistent with this result, neither EdU-positive nor PH3-positive cell density at 72 hpa was significantly altered by transient treatment with these inhibitors (Sup. Fig. S7). At a higher concentration, 10 μM Z-DEVD-FMK produced a modest but statistically significant reduction in tentacle elongation (Fig. 2H, 2I). Together, these results indicate that early caspase-mediated cell death is largely dispensable for blastema formation and regenerative proliferation in *Cladonema* tentacles, although the modest reduction in tentacle elongation observed at a higher concentration of Z-DEVD-FMK raises the possibility that caspase activity may influence later regenerative outgrowth.

### The ERK/MAPK pathway preferentially regulates RSPC-associated blastema formation

In the *Cladonema* medusa tentacle, two distinct undifferentiated proliferative populations contribute to regeneration with different roles: RHSCs predominantly localized in the bulb, which contribute to tissue production during homeostasis and regeneration, including nematocyte formation, and RSPCs, which appear transiently after injury and actively contribute to blastema formation, preferentially differentiating into epithelial cells (Fujita et al., 2023). However, the molecular mechanisms that differentially regulate these two proliferative populations remain unclear. In bilaterians, ERK/MAPK signaling plays an essential role in blastema formation during appendage regeneration, including cockroach leg and zebrafish caudal fin regeneration (Zhang et al., 2024). Given that ERK activation is rapidly induced after amputation and enriched at the injury site in *Cladonema* (Fig. 1E-1G), we hypothesized that ERK/MAPK signaling preferentially regulates blastema formation.

To investigate whether ERK/MAPK signaling regulates blastema formation in *Cladonema*, we inhibited ERK phosphorylation using the MEK inhibitors U0126 and PD184352. As shown above, U0126 significantly reduced injury-induced ERK activation (Fig. 1F, 1G). We further confirmed that PD184352 significantly reduced, but did not completely abolish, the p-ERK signal at 0.5 hpa (Sup. Fig. S5). We then examined cell proliferation by EdU pulse labeling at 24 hpa. U0126 treatment resulted in a concentration-dependent reduction in EdU-positive cell density within the blastema region (Fig. 3A-3C). Similar effects on blastema proliferation were observed with PD184352 treatment (Fig. 3A-3D). At lower inhibitor concentrations, EdU-positive cell density in the bulb remained comparable to controls. At higher concentrations, PD184352 significantly reduced EdU-positive cell density in the bulb, whereas U0126 caused only a modest, non-significant reduction (Fig. 3D). Thus, blastema proliferation was more consistently sensitive to MEK inhibition than proliferation in the bulb, suggesting that ERK/MAPK signaling preferentially promotes the injury-induced RSPC-associated blastema response.

**Figure 3.**
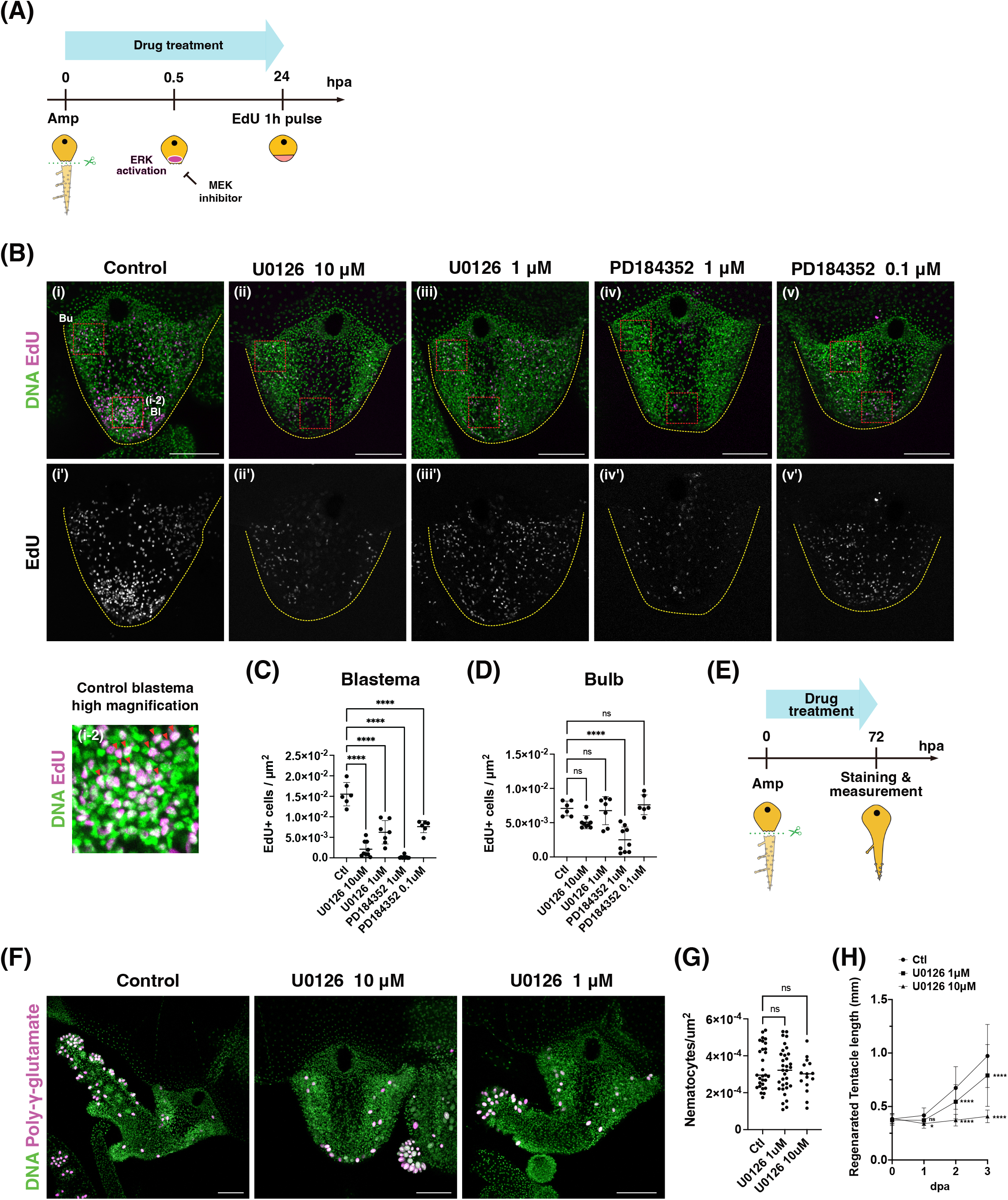
ERK/MAPK signaling preferentially promotes blastema-associated proliferation during tentacle regeneration. **(A)** Experimental scheme for pharmacological inhibition of ERK/MAPK signaling with the MEK inhibitors U0126 or PD184352 during tentacle regeneration. Proliferative activity was assessed by EdU pulse labeling at 24 hpa. **(B)** Detection of S-phase cells by EdU incorporation in regenerating tentacles treated with vehicle control or MEK inhibitors. Representative maximum-intensity projections of z-stacks at 24 hpa are shown. Red dashed boxes indicate the bulb (Bu) and blastema (Bl) regions used for quantification. **(C)** Quantification of EdU-positive cell density within the blastema region. Both U0126 and PD184352 significantly reduced blastema proliferation in a dose-dependent manner. Control: n = 6; U0126, 10 μM: n = 9; U0126, 1 μM: n = 7; PD184352, 1 μM: n = 9; PD184352, 0.1 μM: n = 6. **(D)** Quantification of EdU-positive cell density in the tentacle bulb region. PD184352 at 1 μM significantly reduced EdU-positive cell density, whereas U0126 at 10 μM caused only a modest, non-significant reduction. Lower concentrations of U0126 (1 μM) and PD184352 (0.1 μM) did not significantly affect EdU-positive cell density. Sample numbers are identical to those shown in (C). **(E)** Experimental scheme for assessing nematocyte differentiation and regeneration following MEK inhibition. Animals were treated with U0126 from amputation until 72 hpa, when staining and measurements were performed. **(F)** Poly-γ-glutamate staining of mature nematocytes in regenerating tentacles treated with vehicle control or U0126 (1 or 10 μM) at 72 hpa. Representative maximum-intensity projections of z-stacks are shown. **(G)** Quantification of mature nematocyte density normalized to tentacle area. No significant differences were detected. Control: n = 29; U0126, 10 μM: n = 15; U0126,1 μM: n = 33. **(H)** Quantification of regenerated tentacle length over 72 hpa in control animals and animals treated with 1 or 10 μM U0126. U0126 significantly reduced tentacle elongation in a concentration-dependent manner. Control: n = 54; U0126, 10 μM: n = 36; U0126, 1 μM: n = 54. Data are shown as mean ± SD. Statistical significance was assessed by one-way ANOVA followed by Dunnett’s multiple-comparison test in (C), (D) and (G), and two-way ANOVA followed by Dunnett’s multiple-comparison test in (H). Scale bars: (B, F) 100 μm.

To further examine whether MEK inhibition broadly affects homeostatic proliferative activity in RHSCs, we treated intact tentacles with U0126 for 24 hours and examined EdU incorporation in the bulb. EdU-positive cells were readily detected in the bulb of U0126-treated tentacles, similar to controls (Sup. Fig. S8A-S8C), indicating that U0126 does not detectably impair homeostatic proliferative activity in the bulb. We further performed EdU chase experiments along with U0126 treatment to examine expansion of pre-labeled RHSCs after tentacle amputation under ERK suppression. EdU chase experiments revealed that the labeled RHSC population expanded to a similar extent in both control and U0126-treated animals at 24 hpa (Sup. Fig. S8D-S8F), indicating that U0126 did not detectably alter RHSC expansion during regeneration.

To support the idea that ERK inhibition primarily affects blastema/RSPC dynamics rather than broadly impairing bulb-derived outputs, we assessed nematocyte production by quantifying mature nematocytes using established DAPI-based detection of poly-γ-glutamate (Szczepanek et al., 2002). Although U0126 treatment reduced the total number of nematocytes in a concentration-dependent manner, the density of nematocytes normalized to tentacle area did not significantly differ between treated and control groups (Fig. 3F, 3G). Consistent with enrichment of ERK activation around the injury site (Fig. 1F) and the predominantly epithelial fate of RSPCs (Fujita et al., 2023), these results suggest that ERK inhibition does not strongly impair nematocyte production by RHSCs. Rather, nematocyte number appears to scale with overall tentacle growth, which is reduced upon ERK inhibition.

Given that ERK/MAPK signaling preferentially promotes the blastema response, we next examined whether MEK inhibition affects overall tentacle regeneration. We thus measured the length of regenerating tentacles under U0126 treatment and found that U0126 significantly suppressed regenerative tentacle elongation (Fig. 3H). This phenotype is consistent with previous observations that local UV irradiation, which disrupts blastema formation, inhibits tentacle regeneration (Fujita et al., 2023). Together, these findings indicate that ERK/MAPK signaling preferentially promotes the RSPC-associated blastema response and contributes to efficient tentacle regeneration.

### Notch signaling regulates the balance between nematogenesis and proliferation in RHSCs

While ERK/MAPK signaling preferentially promotes the RSPC-associated blastema response during tentacle regeneration, the mechanisms controlling RHSC behavior are less clear. In cnidarians, including the hydrozoan *Hydra* and *Hydractinia* as well as the anthozoan *Nematostella*, Notch signaling has been implicated in the differentiation of stem/progenitor cells into cnidocytes (Gahan et al., 2017; Käsbauer et al., 2007; Marlow et al., 2012). Because RHSCs, but not RSPCs, contribute to nematocyte production in *Cladonema* tentacles (Fujita et al., 2023), we next examined whether Notch signaling similarly regulates nematocyte formation during regeneration.

To test this, we treated amputated tentacles with the γ-secretase inhibitor DAPT (N-[N-(3,5-difluorophenacetyl)-L-alanyl]-S-phenylglycine tert-butyl ester) or a second γ-secretase inhibitor, LY411575, and examined nematocyte formation during regeneration (Fig. 4A). Both DAPT and LY411575 treatment completely abolished the appearance of mature nematocytes in regenerating tentacles at 3 dpa (Fig. 4B), supporting a requirement for γ-secretase/Notch signaling in nematogenesis in the *Cladonema* medusa tentacle. Both treatments also significantly impaired regenerative tentacle elongation (Fig. 4C), consistent with previous studies where DAPT treatment disrupted tentacle regeneration in *Hydractinia* (Gahan et al., 2017). These results indicate that γ-secretase/Notch signaling is required for nematogenesis during *Cladonema* tentacle regeneration.

**Figure 4.**
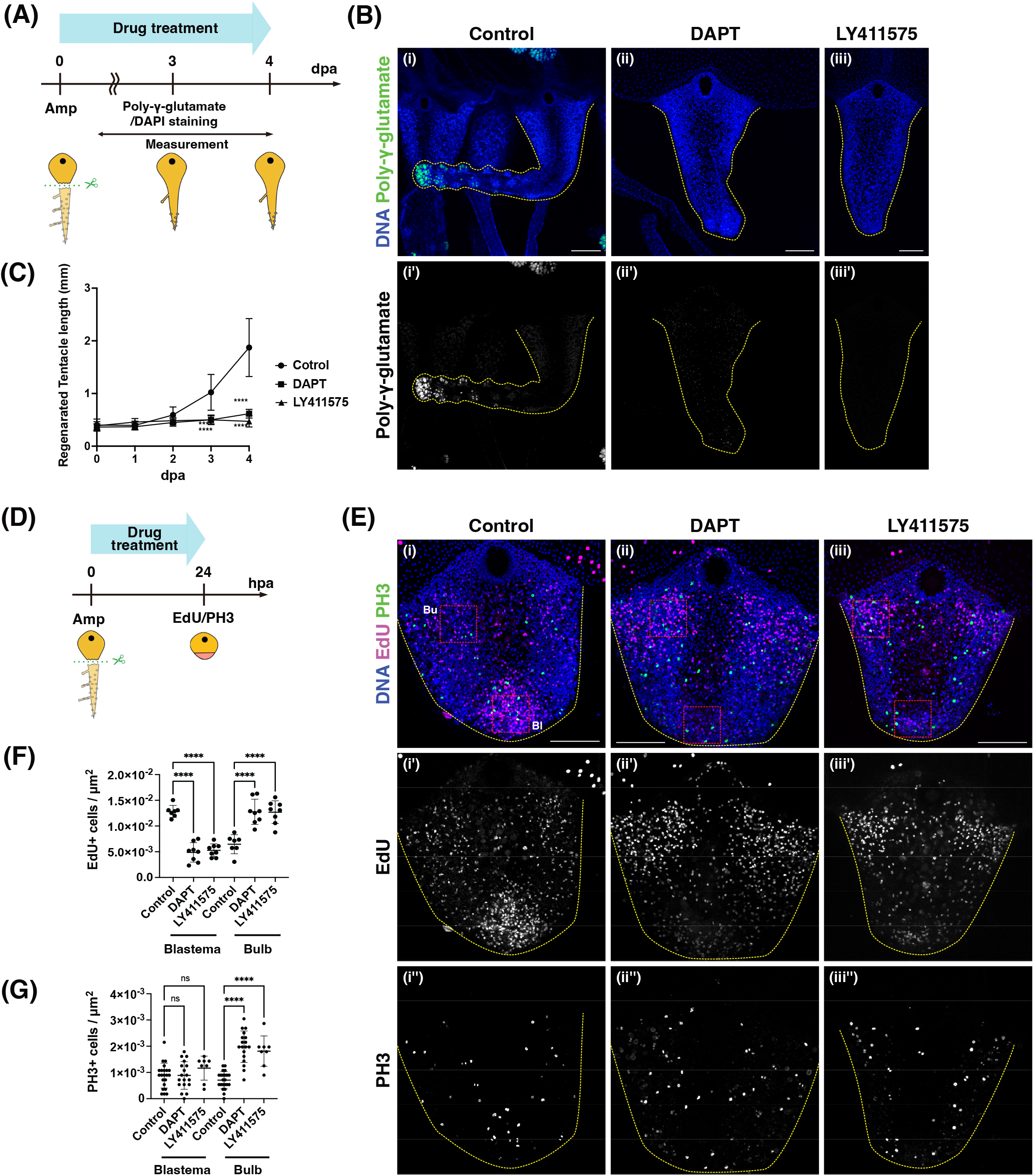
Notch signaling regulates the balance between nematogenesis and proliferation in RHSCs. **(A)** Experimental scheme for pharmacological inhibition of Notch signaling using the γ-secretase inhibitors DAPT (10 μM) and LY411575 (5 μM) during tentacle regeneration. Inhibitor treatment was initiated immediately after amputation and continued until 4 dpa. **(B)** Representative images of regenerating tentacles treated with vehicle control, DAPT or LY411575 at 3 dpa. Mature nematocytes were detected by poly-γ-glutamate staining with DAPI. Both DAPT and LY411575 treatments abolished the appearance of mature nematocytes in regenerating tentacles. **(C)** Quantification of regenerated tentacle length following DAPT and LY411575 treatment. Control: n = 16; DAPT: n = 20; LY411575: n = 10. **(D)** Experimental scheme for assessing proliferative responses under Notch inhibition. **(E)** Detection of S-phase cells by EdU incorporation and M-phase cells by PH3 staining in regenerating tentacles at 24 hpa. Representative maximum-intensity projections of z-stacks are shown. Red dashed boxes indicate the bulb (Bu) and blastema (Bl) regions used for quantification. **(F)** Quantification of EdU-positive cell density in the blastema and bulb regions. Both DAPT and LY411575 reduced EdU-positive cell density in the blastema while increasing EdU-positive cell density in the bulb. Control: n = 7; DAPT: n = 8; LY411575: n = 8. **(G)** Quantification of PH3-positive cell density in the blastema and bulb regions. Both DAPT and LY411575 increased PH3-positive cell density in the bulb, whereas no significant differences were detected in the blastema. Control: n = 22; DAPT: n = 18; LY411575: n = 8. Data are shown as mean ± SD. Statistical significance was assessed by two-way ANOVA followed by Dunnett’s multiple-comparison test in (C) and by one-way ANOVA followed by Šídák’s multiple-comparisons test in (F) and (G). Scale bars: (B), (E) 100 μm.

To determine whether Notch signaling also regulates proliferative activity in *Cladonema*, we examined proliferating cells following γ-secretase inhibition using both EdU incorporation and PH3 staining (Fig. 4D, 4E). In the tentacle bulb, both DAPT and LY411575 significantly increased the density of EdU-positive and PH3-positive cells (Fig. 4F, 4G). In contrast, within the blastema, EdU-positive cell density was significantly reduced by both inhibitors, whereas PH3-positive cell density remained comparable to controls (Fig. 4F, 4G). Together, these results suggest that Notch signaling promotes nematocyte differentiation while restraining proliferative activity in the RHSCs, thereby contributing to the balance between proliferation and differentiation during tentacle regeneration.

## Discussion

In this study, we dissected the early signaling mechanisms underlying tentacle regeneration in the hydrozoan jellyfish *Cladonema*, focusing on the regulation of two distinct undifferentiated proliferative populations: RSPCs and RHSCs (Fig. 5). Our results indicate that tentacle regeneration is not driven by a single, unified stem cell response, but instead relies on pathway-specific control of functionally distinct cell populations. Specifically, ERK/MAPK signaling preferentially regulates the RSPC-associated blastema response at the injury site, whereas Notch signaling predominantly regulates the balance between proliferation and differentiation in RHSCs in the tentacle bulb. In contrast, although extensive cell death occurs immediately after injury, it is largely dispensable for blastema formation and regenerative proliferation. Together, these findings suggest that tentacle regeneration is achieved through differential regulation of distinct proliferative populations, rather than a global activation of stem cell proliferation.

**Figure 5.**
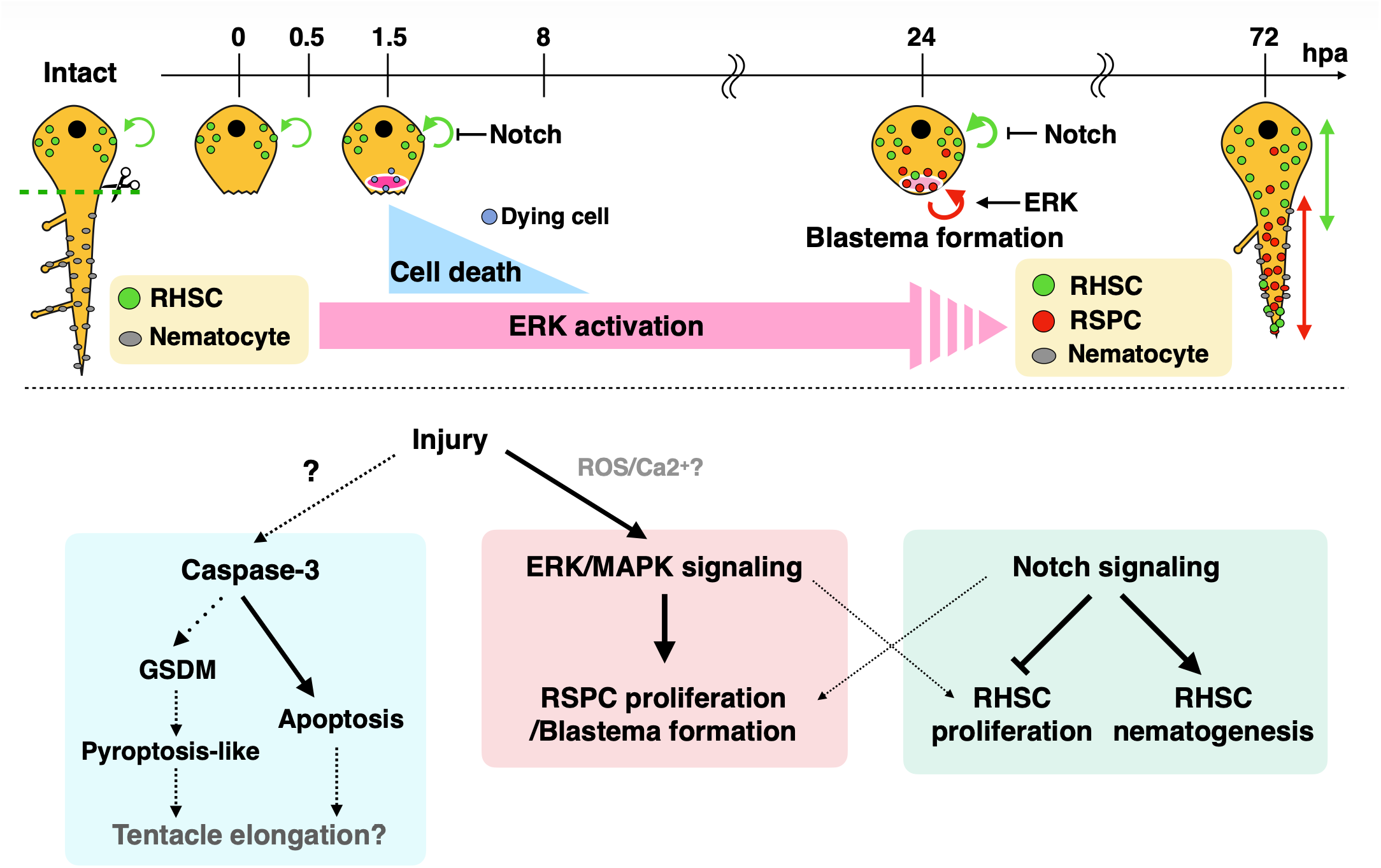
Model for division of labor between proliferative cell populations during *Cladonema* tentacle regeneration. Following tentacle amputation, ERK/MAPK signaling and transient cell death are rapidly induced near the injury site. ERK/MAPK signaling preferentially promotes the RSPC-associated proliferative response and blastema formation. Early caspase-mediated cell death does not provide a major proliferative input to either the RSPC-associated blastema or the RHSC compartment, although a possible contribution to later regenerative outgrowth remains unresolved. In parallel, Notch signaling promotes nematocyte differentiation while restraining proliferative activity in RHSCs. Together, these findings support a model in which tentacle regeneration is achieved through differential deployment of conserved signaling pathways to regulate distinct proliferative cell populations.

### ERK/MAPK signaling preferentially controls blastema formation by RSPCs

The ERK/MAPK signaling pathway is broadly conserved across the animal kingdom and has been implicated in diverse regenerative processes (Wen et al., 2022). In many bilaterian regeneration systems, ERK activation is rapidly induced at wound sites and is required for blastema formation and regenerative outgrowth, including planarian regeneration and zebrafish fin regeneration (Tasaki et al., 2011; Zhang et al., 2024). Our results show that ERK activation is likewise rapidly induced after amputation in *Cladonema* and persists locally at the injury site during blastema formation (Fig. 1E-1G). Importantly, ERK inhibition preferentially impaired proliferative activity within the blastema whereas effects on proliferation in RHSCs were weaker and depended on inhibitor concentration (Fig. 3). U0126 also did not detectably alter homeostatic RHSC proliferation, RHSC expansion during regeneration, or nematocyte density under the conditions examined (Fig. 3; Sup. Fig. S8). Together, these findings suggest that ERK/MAPK signaling preferentially regulates the RSPC-associated blastema response rather than broadly activating proliferative populations throughout the tentacle.

However, our experiments do not establish whether ERK acts cell-autonomously within RSPCs. Because MEK inhibitors were applied systemically and p-ERK-positive cells were not directly identified as RSPCs, ERK activation could occur within RSPCs themselves or in neighboring cells that subsequently provide pro-proliferative signals to the blastema. Distinguishing between these possibilities will require cell-type-resolved analysis of ERK activity and pathway manipulation specifically in RSPCs or surrounding cells.

The evolutionary implications of this preferential ERK deployment are noteworthy. Although components of the ERK/MAPK cascade are conserved across metazoan genomes, its requirement for regeneration is not universal. For example, in the ctenophore *Mnemiopsis leidyi*, pharmacological inhibition of MEK with U0126 did not impair regeneration despite the presence of ERK/MAPK pathway components in its genome, although ERK pathway inhibition was not directly validated (Edgar et al., 2021). Together, these observations suggest that the requirement for ERK/MAPK signaling may vary among regenerative systems during evolution.

### Notch signaling regulates differentiation and proliferative activity in the RHSC compartment

While ERK/MAPK signaling preferentially regulates the RSPC-associated blastema response, Notch signaling plays a distinct and complementary role in regulating RHSC behavior. Inhibition of Notch signaling abolished nematocyte formation during tentacle regeneration (Fig. 4A, 4B), consistent with previous reports in polyp-type cnidarians such as *Hydra, Hydractinia* and *Nematostella*, where Notch signaling is required for cnidocyte differentiation (Gahan et al., 2017; Käsbauer et al., 2007; Marlow et al., 2012). To our knowledge, this provides the first functional evidence in a medusa-stage cnidarian that Notch signaling is essential for nematogenesis during tentacle regeneration (Fig. 4).

The effects of Notch signaling on stem/progenitor proliferation, however, appear to depend on cellular context. During zebrafish fin regeneration, Notch signaling promotes the maintenance and proliferation of blastema progenitor cells, and inhibition of Notch reduces blastema proliferation (Grotek et al., 2013; Münch et al., 2013). In contrast, in *Cladonema*, inhibition of Notch signaling consistently increased both EdU- and PH3-positive cell densities in the RHSC-containing tentacle bulb while impairing nematocyte differentiation. Thus, rather than simply maintaining RHSC-derived cells in a proliferative progenitor state, Notch signaling in the *Cladonema* bulb may promote differentiation and/or restrain proliferative activity. A similar relationship between impaired differentiation and accumulation of proliferative cells has been reported in *Hydra* (Käsbauer et al., 2007) and, more broadly, in the *Drosophila* adult midgut, where loss of Notch blocks differentiation and leads to expansion of proliferative stem/progenitor cells (Micchelli and Perrimon, 2006; Ohlstein and Spradling, 2006).

However, in the blastema, the effect of Notch inhibition was less consistent between cell-cycle markers: PH3-positive cell density was largely unchanged, whereas EdU incorporation was reduced. These results suggest that Notch signaling preferentially regulates proliferative activity and differentiation within the RHSC compartment, while its role in the RSPC-associated blastema response may be more complex and remains to be resolved.

### Early cell death is dispensable for blastema formation in tentacle regeneration

Cell death is frequently observed as an early response to injury during regeneration and has been proposed to promote compensatory proliferation in neighboring cells through the release of mitogenic signals (Bergmann, 2025). Indeed, in cnidarian polyps such as *Hydra*, apoptosis and pyroptosis have been shown to play instructive roles in regeneration by inducing cell proliferation and patterning (Chen et al., 2023a; Chera et al., 2009). In contrast, although we observed extensive early cell death, including an apoptotic component, around the amputation site shortly after tentacle injury (Fig. 1B-1D; Sup. Fig. S3), MEK inhibition did not detectably alter early cell death at 1.5 hpa, suggesting that the early cell-death response is not strongly dependent on ERK/MAPK activity under the conditions examined (Sup. Fig. S6). Conversely, transient inhibition of caspase activity did not significantly affect EdU incorporation or PH3-positive cell density in either the blastema or the tentacle bulb (Fig. 2D-2G). These results indicate that, in contrast to the apoptosis-dependent proliferative response described during *Hydra* head regeneration (Chera et al., 2009; Chera et al., 2011), the early caspase-dependent cell death response is largely dispensable for driving regenerative proliferation in *Cladonema* tentacles and does not appear to provide a major proliferative input to either the RSPC-associated blastema or the RHSC compartment.

Notably, compensatory proliferation driven by cell death has been most clearly demonstrated in tissues in which proliferative stem or progenitor cells are broadly distributed, such as the *Drosophila* wing disc, the mammalian liver, and the *Hydra* body column (Bergantinos et al., 2010; Chera et al., 2009; Fausto, 2004). In contrast, in regenerative systems such as salamander limb regeneration, where blastema formation involves lineage-restricted cells and dedifferentiation rather than uniformly distributed stem cells (Sandoval-Guzman et al., 2014), cell death has not been clearly demonstrated to act as a primary instructive signal. One possible explanation for the limited proliferative response to cell death in *Cladonema* is the spatially compartmentalized organization of its proliferative cell populations, although this possibility remains to be tested experimentally.

Interestingly, at concentrations that were directly shown to suppress early caspase-mediated cell death, transient caspase inhibition did not significantly affect subsequent tentacle elongation. However, a modest reduction in tentacle outgrowth was observed at a higher concentration of Z-DEVD-FMK (Fig. 2H, 2I). Because this effect was observed only at the higher concentration, its mechanistic basis remains uncertain. Previous studies have proposed roles for cell death in tissue remodeling and patterning during regeneration (Pellettieri et al., 2010; Vriz et al., 2014). Thus, although our data do not support a major role for early caspase-mediated cell death in proliferative control, they do not exclude the possibility that caspase-dependent processes contribute to other aspects of regenerative tissue remodeling or patterning (Nakajima and Kuranaga, 2017; Pellettieri et al., 2010; Vriz et al., 2014). These findings highlight potential diversification of regeneration strategies within cnidarians, whereby similar injury responses may be deployed differently across species and regenerative contexts.

### Regeneration as a transient reorganization of cellular society

Our results support a model in which tentacle regeneration is achieved through the differential regulation of distinct proliferative cell populations (Fig. 5). RSPCs are transiently induced at the injury site with ERK/MAPK signaling preferentially promoting the RSPC-associated blastema response (Fig. 3), whereas Notch signaling predominantly regulates proliferation and nematocyte differentiation in the RHSC compartment (Fig. 4), thereby supporting continued production of differentiated cell types during regeneration. Early cell death accompanies injury but does not function as a primary driver of regenerative proliferation (Fig. 2).

This division of labor between conserved signaling pathways may represent a general strategy by which regenerating tissues coordinate distinct cellular populations. Our findings suggest that regeneration is not simply an amplification of homeostatic stem cell activity but instead represents a temporary reorganization of cellular roles within the tissue. Such transient reorganization is consistent with the broader view that regenerative responses involve dynamic changes in cellular states and plasticity (Adler and Sánchez Alvarado, 2015; Nagai and Nakajima, 2026). This modular control of cell populations may allow robust regeneration while preserving tissue integrity and functional specialization (Bangru et al., 2026). Understanding how conserved signaling pathways are differentially deployed to regulate distinct cellular states will be essential for uncovering general principles governing regeneration across animal evolution.

## Declarations

### Ethics approval and consent to participate

Not applicable.

### Consent for publication

Not applicable.

### Availability of data and material

Data will be made available upon reasonable request.

### Authors’ contributions

S.F.: performed most experiments, analyzed data and prepared figures and wrote draft. K.O.: performed experiments and analyzed data. R.K.: performed experiments. R.N.: performed experiments. H.N.: performed experiments. N.S.: analyzed data and edited manuscripts. M.M.: supervised project and edited manuscript. Y.N.: supervised the whole project and experiments and prepared figures and wrote/edited manuscript.

## Acknowledgements

Not applicable.

## Supplementary files

**Figure S1. Conservation of caspase-3 between *Hydra* and *Cladonema*** Pairwise sequence alignment of *Hydra vulgaris* caspase-3B (HyCaspB; AF155128.1) and the predicted *Cladonema pacificum* caspase-3 (CpCaspase-3). The conserved catalytic histidine and cysteine residues are indicated by arrows, and the regions corresponding to the large and small caspase subunits are indicated above the alignment. Strictly identical residues are shown as white letters on a red background, whereas similar residues are shown as red letters; blue frames indicate conserved positions according to the ESPript similarity criteria. The alignment was generated using ClustalW and visualized with ESPript.

**Figure S2. A putative caspase-3 cleavage site in GSDME is conserved between *Hydra* and *Cladonema*** Pairwise sequence alignment of predicted GSDME proteins from *Hydra vulgaris* and *Cladonema pacificum*. The putative caspase-3 cleavage site is highlighted by a green box. Strictly identical residues are shown as white letters on a red background, whereas similar residues are shown as red letters; blue frames indicate conserved positions according to the ESPript similarity criteria. The alignment was generated using ClustalW and visualized with ESPript.

**Figure S3. Apoptotic cell death during the early stage of tentacle regeneration (A)** Detection of apoptotic cell death using TUNEL staining. Representative maximum-intensity projections of z-stacks from intact and regenerating tentacles at 1.5 hpa. Nuclei were counterstained with Hoechst. Red arrowheads indicate TUNEL-positive cells, whereas yellow arrowheads indicate non-specific signals near the eye spot. **(B)** Quantification of TUNEL-positive cells. The number of TUNEL-positive cells was significantly increased at 1.5 hpa compared with intact tentacles. Intact: n = 5; 1.5 hpa: n = 9. Data are shown as mean ± SD. Statistical significance was assessed using two-tailed unpaired t-tests. Scale bars: 100 μm.

**Figure S4. Conservation of the ERK activation-loop TEY motif between *Hydra* and *Cladonema***Pairwise sequence alignment of predicted ERK proteins from *Hydra vulgaris* and *Cladonema pacificum*. The conserved TEY phosphorylation motif within the activation loop is highlighted by a green box. Strictly identical residues are shown as white letters on a red background, whereas similar residues are shown as red letters; blue frames indicate conserved positions according to the ESPript similarity criteria. The alignment was generated using ClustalW and visualized with ESPript.

**Figure S5. PD184352 reduces ERK activation during tentacle regeneration (A)** Immunofluorescence staining for p-ERK in intact tentacles and regenerating tentacles at 0.5 hpa treated with vehicle control or 1 μM PD184352. PD184352 was applied immediately after amputation and maintained until 0.5 hpa. Representative maximum-intensity projections of z-stacks are shown. Nuclei were counterstained with Hoechst. **(B)** Quantification of relative p-ERK fluorescence intensity in the injury-proximal region. PD184352 significantly reduced injury-induced p-ERK intensity compared with controls, although p-ERK remained significantly elevated relative to intact tentacles. Intact: n = 3; control: n = 9; PD184352: n = 9. Data are shown as mean ± SD. Statistical significance was assessed using one-way ANOVA followed by Tukey’s multiple-comparison test. Scale bars: 100 μm.

**Figure S6. MEK inhibition does not detectably alter early injury-induced cell death (A)** SYTOX Green staining of regenerating tentacles at 1.5 hpa treated with vehicle control, 10 μM U0126, or 1 μM PD184352. MEK inhibitors were applied immediately after amputation and maintained until 1.5 hpa. Representative maximum-intensity projections of z-stacks are shown. Nuclei were counterstained with Hoechst. **(B)** Quantification of background-corrected SYTOX Green fluorescence intensity in the injury-proximal region. Neither U0126 nor PD184352 significantly altered SYTOX fluorescence intensity compared with controls. **(C)** Quantification of SYTOX-positive cell density within the same region. Only SYTOX signals overlapping Hoechst-positive nuclei were counted, and obvious non-specific fluorescence was excluded from the analysis. Neither U0126 nor PD184352 significantly altered SYTOX-positive cell density. Control: n = 9; U0126: n = 7; PD184352: n = 8. Data are shown as mean ± SD. Statistical significance was assessed by one-way ANOVA followed by Dunnett’s multiple-comparison test in (B) and one-way ANOVA followed by Tukey’s multiple-comparison test in (C). Scale bars: 100 μm.

**Figure S7. Transient inhibition of early caspase-mediated cell death does not detectably affect proliferative activity at 72 hpa. (A)** Detection of S-phase and mitotic cells by 1-h EdU incorporation and PH3 immunostaining, respectively, in regenerating tentacles at 72 hpa following transient treatment with 10 μM Q-VD-Oph or 1 μM Z-DEVD-FMK. Caspase inhibitors were applied from 1.5 h before amputation until 1.5 hpa and were then removed. Representative maximum-intensity projections of z-stacks are shown. Nuclei were counterstained with Hoechst. **(B)** Quantification of EdU-positive cell density at 72 hpa. Control: n = 10; Q-VD-Oph: n = 8; Z-DEVD-FMK: n = 11. **(C)** Quantification of PH3-positive cell density at 72 hpa. Control: n = 10; Q-VD-Oph: n = 8; Z-DEVD-FMK: n = 11. Data are shown as mean ± SD. Statistical significance was assessed by one-way ANOVA followed by Dunnett’s multiple-comparisons test. No significant differences were detected between control and inhibitor-treated groups. Scale bar: 100 μm.

**Figure S8. U0126 does not detectably alter RHSC proliferation or expansion (A)** Experimental scheme for drug treatment of intact tentacles and detection of proliferating cells by EdU pulse labeling. Intact tentacles were treated for 24 h without amputation, followed by a 1-h EdU pulse labeling. **(B)** Detection of EdU-positive cells in intact tentacles treated with vehicle control, U0126 (1 or 10 μM), or hydroxyurea (HU; 10 mM), which was used as a positive control for inhibition of cell proliferation. Representative maximum-intensity projections of z-stacks are shown. **(C)** Quantification of EdU-positive cell density in the tentacle bulb region. Control: n = 13; U0126, 1 μM: n = 12; U0126, 10 μM: n = 9; HU: n = 7. **(D)** Experimental scheme of EdU chase analysis of pre-existing RHSCs shown in (E). EdU pulse labeling was performed for 1 h before amputation, and U0126 treatment was initiated immediately after amputation and maintained until 24 hpa, when EdU-positive cells were analyzed. **(E)** Detection of EdU-labeled pre-existing RHSCs in regenerating tentacles treated with vehicle control or U0126 (1 or 10 μM). Representative maximum-intensity projections of z-stacks at 24 hpa are shown. **(F)** Quantification of EdU-positive cell density in the tentacle bulb during regeneration. Before amputation: n = 11; Control: n = 11; U0126, 1 μM: n = 11; U0126, 10 μM: n = 11. Data are shown as mean ± SD. Statistical significance was assessed by one-way ANOVA followed by Dunnett’s multiple-comparison test in (C) and one-way ANOVA followed by Tukey’s multiple-comparison test in (F). Scale bars: 100 μm.

## References

Accorsi, A., Guo, L., Marshall, W. F., Mommersteeg, M. T. M. and Nakajima, Y. I. (2024). Extraordinary model systems for regeneration. Development 151. 10.1242/dev.203083

Adell, T., Cebria, F., Abril, J. F., Araujo, S. J., Corominas, M., Morey, M., Serras, F. and Gonzalez-Estevez, C. (2025). Cell death in regeneration and cell turnover: Lessons from planarians and Drosophila. Semin Cell Dev Biol 169, 103605. 10.1016/j.semcdb.2025.103605

Adler, C. E. and Sánchez Alvarado, A. (2015). Types or States? Cellular Dynamics and Regenerative Potential. Trends Cell Biol 25, 687–696. 10.1016/j.tcb.2015.07.008

Aztekin, C. and Storer, M. A. (2022). To regenerate or not to regenerate: Vertebrate model organisms of regeneration-competency and -incompetency. Wound Repair Regen 30, 623–635. 10.1111/wrr.13000

Bangru, S., Diegmiller, R., Di Talia, S. and Poss, K. D. (2026). Signal control during tissue regeneration in adult animals. Nat Rev Mol Cell Biol 27, 316–335. 10.1038/s41580-025-00917-1

Bely, A. E. and Nyberg, K. G. (2010). Evolution of animal regeneration: re-emergence of a field. Trends Ecol Evol 25, 161–170. 10.1016/j.tree.2009.08.005

Bergantinos, C., Vilana, X., Corominas, M. and Serras, F. (2010). Imaginal discs: Renaissance of a model for regenerative biology. Bioessays 32, 207–217. 10.1002/bies.200900105

Bergmann, A. (2025). Cell Death, Compensatory Proliferation, and Cell Competition. Annu Rev Genet 59, 165–187. 10.1146/annurev-genet-012125-083359

Chen, S., Gong, Y., Li, S., Yang, D., Zhang, Y. and Liu, Q. (2023a). Hydra gasdermin-gated pyroptosis signalling regulates tissue regeneration. Dev Comp Immunol 149, 104904. 10.1016/j.dci.2023.104904

Chen, S., Li, S., Chen, H., Gong, Y., Yang, D., Zhang, Y. and Liu, Q. (2023b). Caspase-mediated LPS sensing and pyroptosis signaling in Hydra. Sci Adv 9, eadh4054. 10.1126/sciadv.adh4054

Chera, S., Ghila, L., Dobretz, K., Wenger, Y., Bauer, C., Buzgariu, W., Martinou, J. C. and Galliot, B. (2009). Apoptotic cells provide an unexpected source of Wnt3 signaling to drive hydra head regeneration. Dev Cell 17, 279–289. 10.1016/j.devcel.2009.07.014

Chera, S., Ghila, L., Wenger, Y. and Galliot, B. (2011). Injury-induced activation of the MAPK/CREB pathway triggers apoptosis-induced compensatory proliferation in hydra head regeneration. Dev Growth Differ 53, 186–201. 10.1111/j.1440-169X.2011.01250.x

Denker, E., Manuel, M., Leclere, L., Le Guyader, H. and Rabet, N. (2008). Ordered progression of nematogenesis from stem cells through differentiation stages in the tentacle bulb of Clytia hemisphaerica (Hydrozoa, Cnidaria). Dev Biol 315, 99–113. 10.1016/j.ydbio.2007.12.023

DuBuc, T. Q., Traylor-Knowles, N. and Martindale, M. Q. (2014). Initiating a regenerative response; cellular and molecular features of wound healing in the cnidarian Nematostella vectensis. BMC Biol 12, 24. 10.1186/1741-7007-12-24

Edgar, A., Mitchell, D. G. and Martindale, M. Q. (2021). Whole-Body Regeneration in the Lobate Ctenophore Mnemiopsis leidyi. Genes (Basel) 12. 10.3390/genes12060867

Fausto, N. (2004). Liver regeneration and repair: hepatocytes, progenitor cells, and stem cells. Hepatology 39, 1477–1487. 10.1002/hep.20214

Fujita, S., Kuranaga, E., Miura, M. and Nakajima, Y. I. (2022). Fluorescent In Situ Hybridization and 5-Ethynyl-2’-Deoxyuridine Labeling for Stem-like Cells in the Hydrozoan Jellyfish Cladonema pacificum. J Vis Exp. 10.3791/64285

Fujita, S., Kuranaga, E. and Nakajima, Y. I. (2019). Cell proliferation controls body size growth, tentacle morphogenesis, and regeneration in hydrozoan jellyfish Cladonema pacificum. PeerJ 7, e7579. 10.7717/peerj.7579

Fujita, S., Kuranaga, E. and Nakajima, Y. I. (2021). Regeneration Potential of Jellyfish: Cellular Mechanisms and Molecular Insights. Genes (Basel) 12. 10.3390/genes12050758

Fujita, S., Takahashi, M., Kumano, G., Kuranaga, E., Miura, M. and Nakajima, Y. I. (2023). Distinct stem-like cell populations facilitate functional regeneration of the Cladonema medusa tentacle. PLoS Biol 21, e3002435. 10.1371/journal.pbio.3002435

Gahan, J. M., Bradshaw, B., Flici, H. and Frank, U. (2016). The interstitial stem cells in Hydractinia and their role in regeneration. Curr Opin Genet Dev 40, 65–73. 10.1016/j.gde.2016.06.006

Gahan, J. M., Schnitzler, C. E., DuBuc, T. Q., Doonan, L. B., Kanska, J., Gornik, S. G., Barreira, S., Thompson, K., Schiffer, P., Baxevanis, A. D., et al. (2017). Functional studies on the role of Notch signaling in Hydractinia development. Dev Biol 428, 224–231. 10.1016/j.ydbio.2017.06.006

Gauron, C., Rampon, C., Bouzaffour, M., Ipendey, E., Teillon, J., Volovitch, M. and Vriz, S. (2013). Sustained production of ROS triggers compensatory proliferation and is required for regeneration to proceed. Sci Rep 3, 2084. 10.1038/srep02084

Grotek, B., Wehner, D. and Weidinger, G. (2013). Notch signaling coordinates cellular proliferation with differentiation during zebrafish fin regeneration. Development 140, 1412–1423. 10.1242/dev.087452

Guerin, D. J., Kha, C. X. and Tseng, K. A. (2021). From Cell Death to Regeneration: Rebuilding After Injury. Front Cell Dev Biol 9, 655048. 10.3389/fcell.2021.655048

Hobmayer, B., Rentzsch, F., Kuhn, K., Happel, C. M., von Laue, C. C., Snyder, P., Rothbacher, U. and Holstein, T. W. (2000). WNT signalling molecules act in axis formation in the diploblastic metazoan Hydra. Nature 407, 186–189. 10.1038/35025063

Holstein, T. W., Hobmayer, E. and Technau, U. (2003). Cnidarians: an evolutionarily conserved model system for regeneration? Dev Dyn 226, 257–267. 10.1002/dvdy.10227

Juanico, I. Y., Stockinger, A. W., Virgen, A. K., Srisrimal, N., Campos, S. E. and Juliano, C. E. (2026). Sustained ERK signaling couples the injury response to organizer formation during Hydra head regeneration. Dev Biol 535, 42–57. 10.1016/j.ydbio.2026.03.018

Kanehisa, R., Nakatani, H., Takatori, S., Tomita, T., Miura, M. and Nakajima, Y. I. (2026). Aging Disrupts Tissue Homeostasis and Constrains Blastema-Mediated Regeneration in the Cladonema Medusa. Aging Cell 25, e70727. 10.1111/acel.70727

Kawamoto, Y., Nakajima, Y. I. and Kuranaga, E. (2016). Apoptosis in Cellular Society: Communication between Apoptotic Cells and Their Neighbors. Int J Mol Sci 17, 2144. 10.3390/ijms17122144

Kragl, M., Knapp, D., Nacu, E., Khattak, S., Maden, M., Epperlein, H. H. and Tanaka, E. M. (2009). Cells keep a memory of their tissue origin during axolotl limb regeneration. Nature 460, 60–65. 10.1038/nature08152

Käsbauer, T., Towb, P., Alexandrova, O., David, C. N., Dall’armi, E., Staudigl, A., Stiening, B. and Bottger, A. (2007). The Notch signaling pathway in the cnidarian Hydra. Dev Biol 303, 376–390. 10.1016/j.ydbio.2006.11.022

Leclère, L., Jager, M., Barreau, C., Chang, P., Le Guyader, H., Manuel, M. and Houliston, E. (2012). Maternally localized germ plasm mRNAs and germ cell/stem cell formation in the cnidarian Clytia. Dev Biol 364, 236–248. 10.1016/j.ydbio.2012.01.018

Lee, Y., Grill, S., Sanchez, A., Murphy-Ryan, M. and Poss, K. D. (2005). Fgf signaling instructs position-dependent growth rate during zebrafish fin regeneration. Development 132, 5173–5183. 10.1242/dev.02101

Macias-Munoz, A. (2025). Investigating the evolution and features of regeneration using cnidarians. Integr Comp Biol 65, 713–726. 10.1093/icb/icaf006

Marlow, H., Roettinger, E., Boekhout, M. and Martindale, M. Q. (2012). Functional roles of Notch signaling in the cnidarian Nematostella vectensis. Dev Biol 362, 295–308. 10.1016/j.ydbio.2011.11.012

Masuda-Ozawa, T., Fujita, S., Nakamura, R., Watanabe, H., Kuranaga, E. and Nakajima, Y. I. (2022). siRNA-mediated gene knockdown via electroporation in hydrozoan jellyfish embryos. Sci Rep 12, 16049. 10.1038/s41598-022-20476-1

Micchelli, C. A. and Perrimon, N. (2006). Evidence that stem cells reside in the adult Drosophila midgut epithelium. Nature 439, 475–479. 10.1038/nature04371

Min, S. and Whited, J. L. (2023). Limb blastema formation: How much do we know at a genetic and epigenetic level? J Biol Chem 299, 102858. 10.1016/j.jbc.2022.102858

Münch, J., Gonzalez-Rajal, A. and de la Pompa, J. L. (2013). Notch regulates blastema proliferation and prevents differentiation during adult zebrafish fin regeneration. Development 140, 1402–1411. 10.1242/dev.087346

Nagai, H. and Nakajima, Y. I. (2026). Epithelial cell plasticity in metazoans: Evolutionary insights into roles and mechanisms. Semin Cell Dev Biol 179-180, 103670. 10.1016/j.semcdb.2026.103670

Nakajima, Y. I. and Kuranaga, E. (2017). Caspase-dependent non-apoptotic processes in development. Cell Death Differ 24, 1422–1430. 10.1038/cdd.2017.36

Ohlstein, B. and Spradling, A. (2006). The adult Drosophila posterior midgut is maintained by pluripotent stem cells. Nature 439, 470–474. 10.1038/nature04333

Pellettieri, J., Fitzgerald, P., Watanabe, S., Mancuso, J., Green, D. R. and Sanchez Alvarado, A. (2010). Cell death and tissue remodeling in planarian regeneration. Dev Biol 338, 76–85. 10.1016/j.ydbio.2009.09.015

Salvetti, A. and Rossi, L. (2019). Planarian Stem Cell Heterogeneity. Adv Exp Med Biol 1123, 39–54. 10.1007/978-3-030-11096-3_4

Sandoval-Guzman, T., Wang, H., Khattak, S., Schuez, M., Roensch, K., Nacu, E., Tazaki, A., Joven, A., Tanaka, E. M. and Simon, A. (2014). Fundamental differences in dedifferentiation and stem cell recruitment during skeletal muscle regeneration in two salamander species. Cell Stem Cell 14, 174–187. 10.1016/j.stem.2013.11.007

Scimone, M. L., Kravarik, K. M., Lapan, S. W. and Reddien, P. W. (2014). Neoblast specialization in regeneration of the planarian Schmidtea mediterranea. Stem Cell Reports 3, 339–352. 10.1016/j.stemcr.2014.06.001

Sinigaglia, C., Peron, S., Eichelbrenner, J., Chevalier, S., Steger, J., Barreau, C., Houliston, E. and Leclere, L. (2020). Pattern regulation in a regenerating jellyfish. Elife 9. 10.7554/eLife.54868

Srivastava, M. (2021). Beyond Casual Resemblance: Rigorous Frameworks for Comparing Regeneration Across Species. Annu Rev Cell Dev Biol 37, 415–440. 10.1146/annurev-cellbio-120319-114716

Szczepanek, S., Cikala, M. and David, C. N. (2002). Poly-gamma-glutamate synthesis during formation of nematocyst capsules in Hydra. J Cell Sci 115, 745–751. 10.1242/jcs.115.4.745

Takeda, N., Kon, Y., Quiroga Artigas, G., Lapebie, P., Barreau, C., Koizumi, O., Kishimoto, T., Tachibana, K., Houliston, E. and Deguchi, R. (2018). Identification of jellyfish neuropeptides that act directly as oocyte maturation-inducing hormones. Development 145. 10.1242/dev.156786

Tasaki, J., Shibata, N., Nishimura, O., Itomi, K., Tabata, Y., Son, F., Suzuki, N., Araki, R., Abe, M., Agata, K., et al. (2011). ERK signaling controls blastema cell differentiation during planarian regeneration. Development 138, 2417–2427. 10.1242/dev.060764

Tursch, A., Bartsch, N., Mercker, M., Schluter, J., Lommel, M., Marciniak-Czochra, A., Ozbek, S. and Holstein, T. W. (2022). Injury-induced MAPK activation triggers body axis formation in Hydra by default Wnt signaling. Proc Natl Acad Sci U S A 119, e2204122119. 10.1073/pnas.2204122119

van Wolfswinkel, J. C., Wagner, D. E. and Reddien, P. W. (2014). Single-cell analysis reveals functionally distinct classes within the planarian stem cell compartment. Cell Stem Cell 15, 326–339. 10.1016/j.stem.2014.06.007

Vogg, M. C., Galliot, B. and Tsiairis, C. D. (2019). Model systems for regeneration: Hydra. Development 146. 10.1242/dev.177212

Vriz, S., Reiter, S. and Galliot, B. (2014). Cell death: a program to regenerate. Curr Top Dev Biol 108, 121–151. 10.1016/B978-0-12-391498-9.00002-4

Wen, X., Jiao, L. and Tan, H. (2022). MAPK/ERK Pathway as a Central Regulator in Vertebrate Organ Regeneration. Int J Mol Sci 23. 10.3390/ijms23031464

Zeng, A., Li, H., Guo, L., Gao, X., McKinney, S., Wang, Y., Yu, Z., Park, J., Semerad, C., Ross, E., et al. (2018). Prospectively Isolated Tetraspanin+ Neoblasts Are Adult Pluripotent Stem Cells Underlying Planaria Regeneration. Cell 173, 1593–1608.e20. 10.1016/j.cell.2018.05.006

Zhang, X. S., Wei, L., Zhang, W., Zhang, F. X., Li, L., Li, L., Wen, Y., Zhang, J. H., Liu, S., Yuan, D., et al. (2024). ERK-activated CK-2 triggers blastema formation during appendage regeneration. Sci Adv 10, eadk8331. 10.1126/sciadv.adk8331

