## Supplementary Figures for "Division of labor between ERK/MAPK and Notch signaling in regulating distinct proliferative populations during jellyfish tentacle regeneration"

### Figure S1

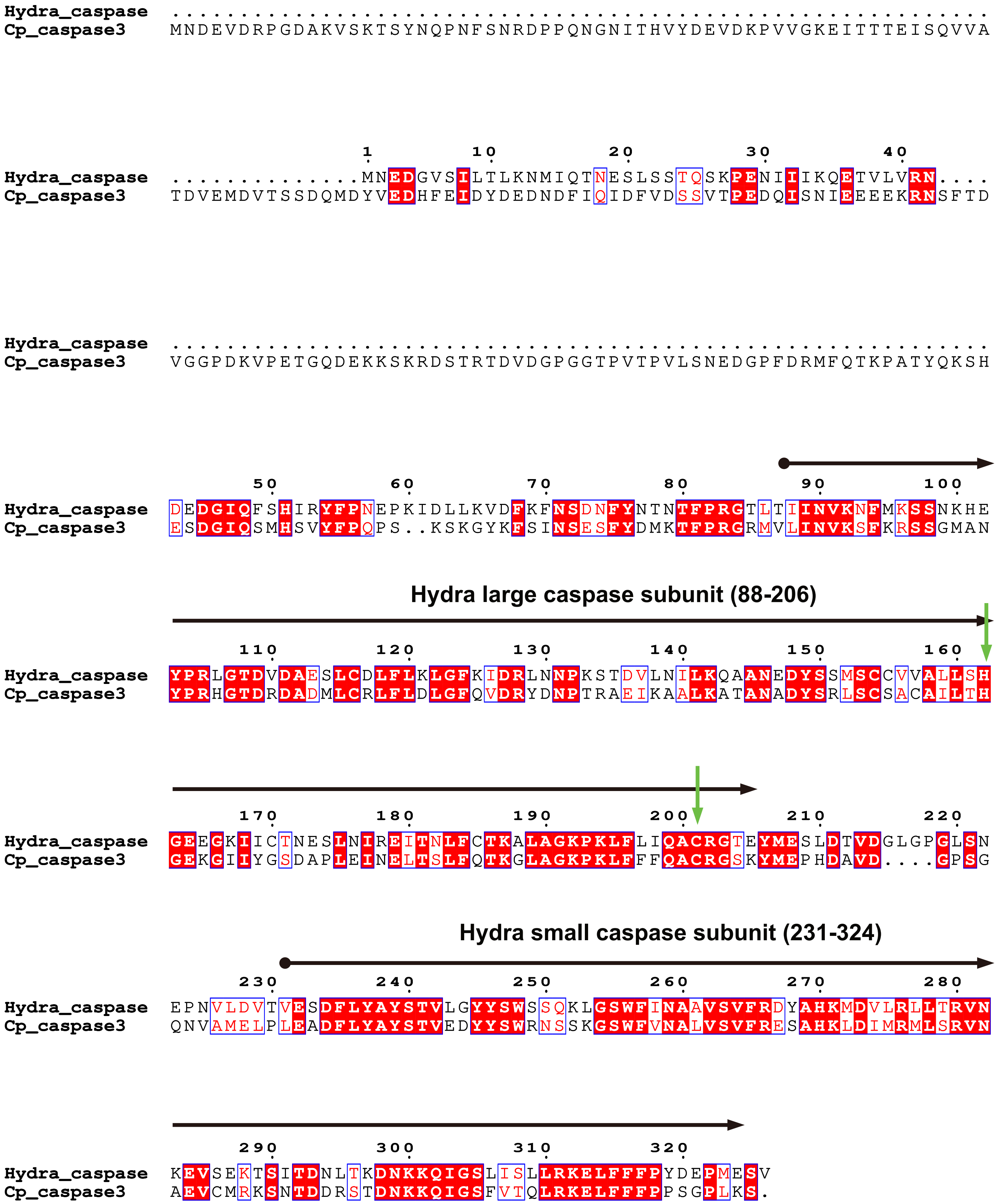
Hydra_caspase ............................................................
Cp_caspase3 MNDEVDRPGDAKVSKTSYNQPNFSNRDPPQNGNITHVYDEVDKPVVGKEITTTEISQVVA
1 10 20 30 40
Hydra_caspase ..............MNEDGVSILTLKNMIQTNESLSSTQSKPENIIIKQETVLVRN....
Cp_caspase3 TDVEMDVTSSDQMDYVEDHFEIDYDEDNDFIQIDFVDSSVTPEDQISNIEEEEKRNSFTD
Hydra_caspase ............................................................
Cp_caspase3 VGGPDKVPETGQDEKKSKRDSTRTDVDGPGGTPVTPVLSNEDGPFDRMFQTKPATYQKSH
50 60 70 80 90 100
Hydra_caspase DEDGIQFSHIRYFPNEPKIDLLKVDFKFNSDNFYNTNTFPRGTLTIINVKNFMKSSNKHE
Cp_caspase3 ESDGIQSMHSVYFPQPS..KSKGYKFSINSESFYDMKTFPRGRMVLINVKSFKRSSGMAN
Hydra large caspase subunit (88-206)
110 120 130 140 150 160
Hydra_caspase YPRLGTDVDAESLCDLFLKLGFKIDRLNNPKSTDVLNILKQAANEDYSSMSCCVVALLSH
Cp_caspase3 YPRHGTDRDADMLCRLFLDLGFQVDRYDNPTRAEIKAALKATANADYSRLSCSACAILTH
170 180 190 200 210 220
Hydra_caspase GEEGKIICTNESLNIREITNLFCTKALAGKPKLFLIQACRGTEYMESLDTVDGLGPGLSN
Cp_caspase3 GEKGIIYGSDAPLEINELTSLFQTKGLAGKPKLFFFQACRGSKYMEPHDAVD....GPSG
Hydra small caspase subunit (231-324)
230 240 250 260 270 280
Hydra_caspase EPNVLDVTVESDFLYAYSTVLGYYSWSSQKLGSWFINAAVSVFRDYAHKMDVLRLLTRVN
Cp_caspase3 QNVAMELPLEADFLYAYSTVEDYYSWRNSSKGSWFVNALVSVFRESAHKLDIMRMLSRVN
290 300 310 320
Hydra_caspase KEVSEKTSITDNLTKDNKKQIGSLISLLRKELFFFPYDEPMESV
Cp_caspase3 AEVCMRKSNTDDRSTDNKKQIGSFVTQLRKELFFFPPSGPLKS.

### Figure S2

```
              1        10        20        30        40        50        60
Hydra_GSDME   MAMFEAATKCIAKSIGSKTLHHVSDLNSSERFKLLCVVCQKKSFWPWKKTQIFPTQIVLQ
Cp_GSDME-like MSMFKEASKHVSRSAGSKTLHPVADLNSSEWYRTLRIVKRRTSFWFWKKAKHIPTQVGLE
```

```
                       70        80        90       100       110
Hydra_GSDME   DILLTKLNIEQSDVN.FTLVKDYIENPTLNIKGDVGAKIASELQLEFSASDSVSLNINVG
Cp_GSDME-like DILTQKLNLDESKFKSSVLLEQFVEKPTFKLQGDIGAKIASELNLDLSASDELHLTLKLG
```

```
              120       130       140       150       160       170
Hydra_GSDME   SLQKQEIKWQKLDNALQSSKVNTRHPLLEEIQFLKRTELAVVLESISTCADSTLSGSADL
Cp_GSDME-like NILKTRVNWEMLQERLLVNSVNLSHRLFKEVQKESRTTLCVVLESLSTEGEGEFEETKDV
```

```
              180           190       200       210       220       230
Hydra_GSDME   NVTTDDSVSN....KSVVVVDVHTKDSIEKKTTHSYTLPSNTVLAFSCNTFSVTEYGGID
Cp_GSDME-like TVSTTDEGTIPKIPKSTITVDVHTKGSVTSETVHSFNIPPKTVLSYSCNEISIDKNGFIS
```

```
                  240       250         260       270       280       290
Hydra_GSDME   FHAAPDEADFSTKTLLQKTK..KDSFDFVKSMLFSFVKSSKFSSIQQFFKSIIGSVEQSS
Cp_GSDME-like LHSTVDSYDSNTEFLILKEVDNENSVESVKRQFIPLLKSEKFDQIKLAFKIFSESANEDS
```

```
                   300           310       320       330       340
Hydra_GSDME   LEAIYLLLLAVE....GSSENNFYLNSCHSALSKTIVENIGLWKPLLLSLNFFIPDDLSL
Cp_GSDME-like LQALHFLLSAAESCLDGRSEDKTFKLSTIKHLMSSCVDES..WGSILKLLGFLLPTNIEN
```

```
              350       360       370       380       390       400
Hydra_GSDME   NKASIKFPTNDDDYLVKGCIGWIDAILDTDKNLRHYLVQFSDEHLKLLLHIVESKILNKE
Cp_GSDME-like ENAEIILPTEDKDNILKSCVAFLDSLEDIEEASRKLLSGIPSEHIKVLMKIVENELHGKE
```

```
              410        420       430       440       450       460
Hydra_GSDME   IN.EEIQINLMFSENQSTKLFLNGIGFVFETEEEKEVMTYPSEDHQKLLDAFINIYLLSA
Cp_GSDME-like NKTKDPLIQNTFLKEPSTRLFLKNVGFIEMCADDEELLSFSSHHSVSLLDIYTVLYVFGS
```

### Figure S3

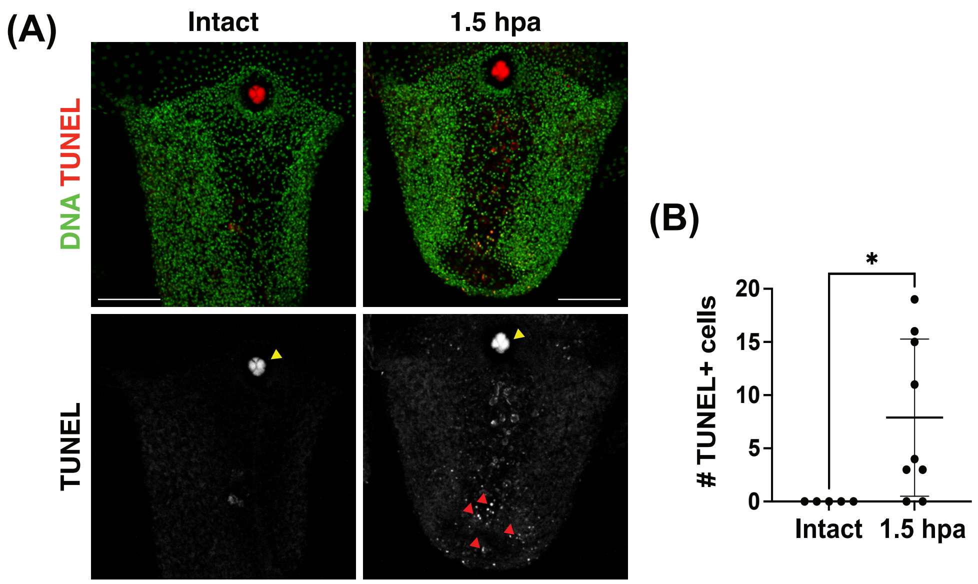

(A)
Intact
1.5 hpa
DNA TUNEL
TUNEL
(B)
*
### TUNEL+ cells
20
15
10
5
0
Intact
1.5 hpa

#### Figure S4

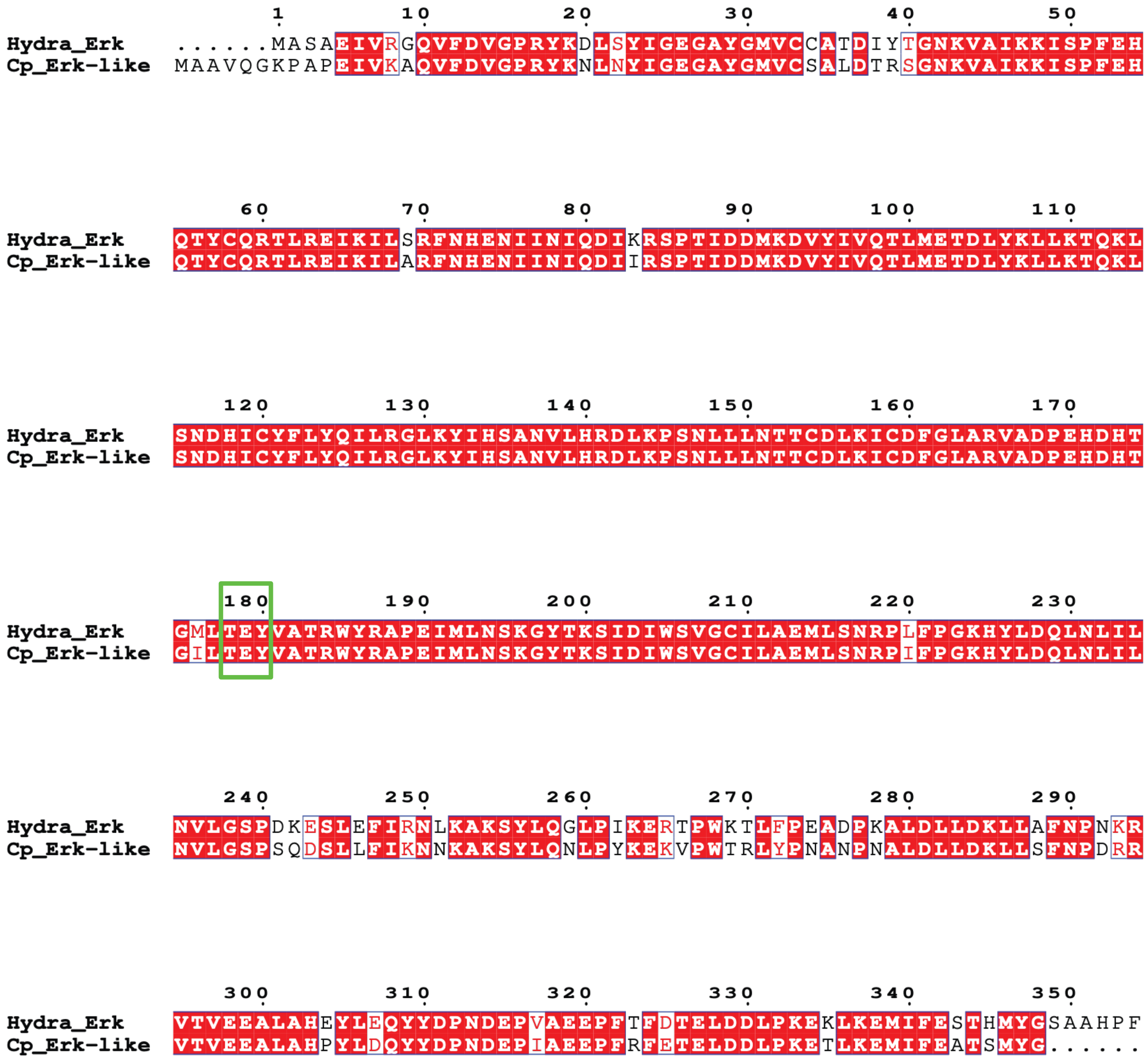
1 10 20 30 40 50
Hydra_Erk ......MASAEIVRGQVFDVGPRYKDLSYIGEGAYGMVCCATDIYTGNKVAIKKISPFEH
Cp_Erk-like MAAVQGKPAPEIVKAQVFDVGPRYKNLNYIGEGAYGMVCSALDTRSGNKVAIKKISPFEH
60 70 80 90 100 110
Hydra_Erk QTYCQRTLREIKILSRFNHENIINIQDIKRSPTIDDMKDVYIVQTLMETDLYKLLKTQKL
Cp_Erk-like QTYCQRTLREIKILARFNHENIINIQDIIRSPTIDDMKDVYIVQTLMETDLYKLLKTQKL
120 130 140 150 160 170
Hydra_Erk SNDHICYFLYQILRGLKYIHSANVLHRDLKPSNLLLNTTCDLKICDFGLARVADPEHDHT
Cp_Erk-like SNDHICYFLYQILRGLKYIHSANVLHRDLKPSNLLLNTTCDLKICDFGLARVADPEHDHT
180 190 200 210 220 230
Hydra_Erk GMLTEYVATRWYRAPEIMLNSKGYTKSIDIWSVGCILAEMLSNRPLFPGKHYLDQLNLIL
Cp_Erk-like GILTEYVATRWYRAPEIMLNSKGYTKSIDIWSVGCILAEMLSNRPIFPGKHYLDQLNLIL
240 250 260 270 280 290
Hydra_Erk NVLGSPDKESLEFIRNLKAKSYLQGLPIKERTPWKTLFPEADPKALDLLDKLLAFNPNKR
Cp_Erk-like NVLGSPSQDSLLFIKNNKAKSYLQNLPYKEKVPWTRLYPNANPNALDLLDKLLSFNPDRR
300 310 320 330 340 350
Hydra_Erk VTVEEALAHEYLEQYYDPNDEPVAEEPFTFDTELDDLPKEKLKEMIFESTHMYGSAAHPF
Cp_Erk-like VTVEEALAHPYLDQYYDPNDEPIAEEPFRFETELDDLPKETLKEMIFEATSMYG......

### Figure S5

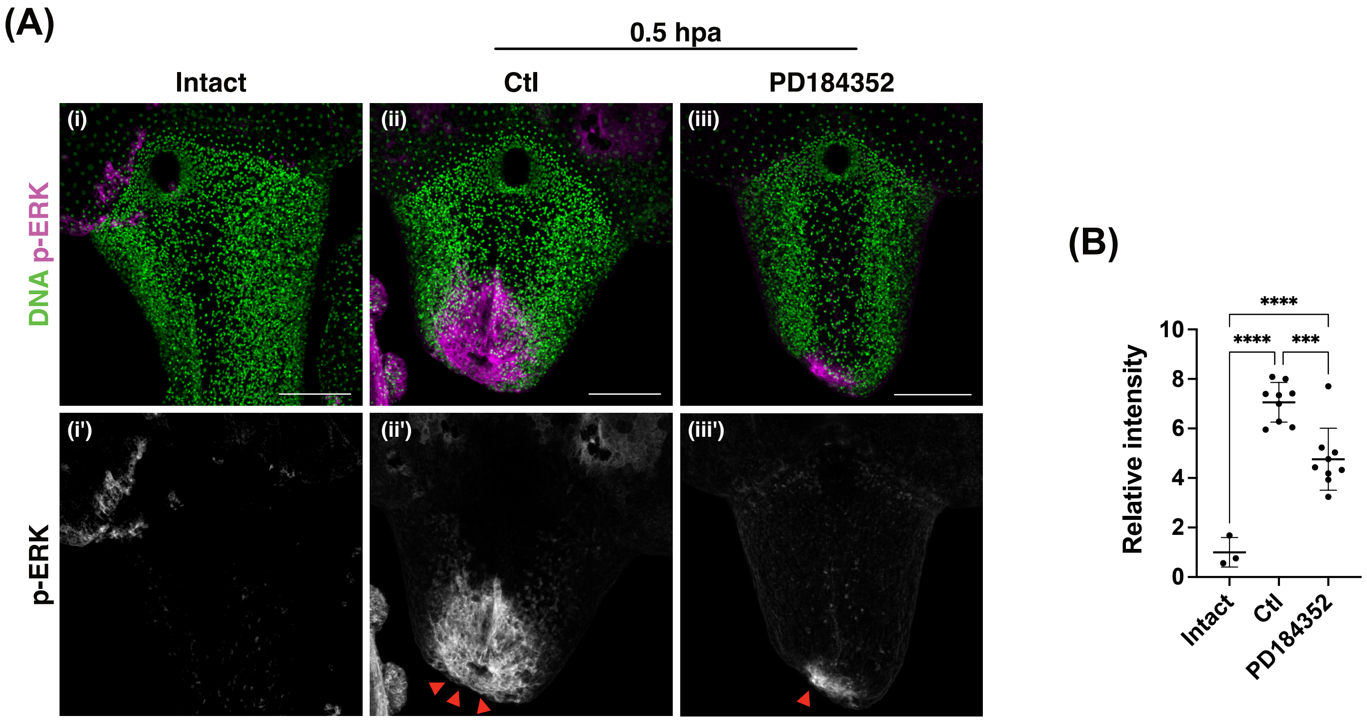
(A)
0.5 hpa
Intact
Ctl
PD184352
(i)
(ii)
(iii)
(i')
(ii')
(iii')
DNA p-ERK
p-ERK
(B)
Relative intensity
0
2
4
6
8
10
****
****
***
Intact
Ctl
PD184352

### Figure S6

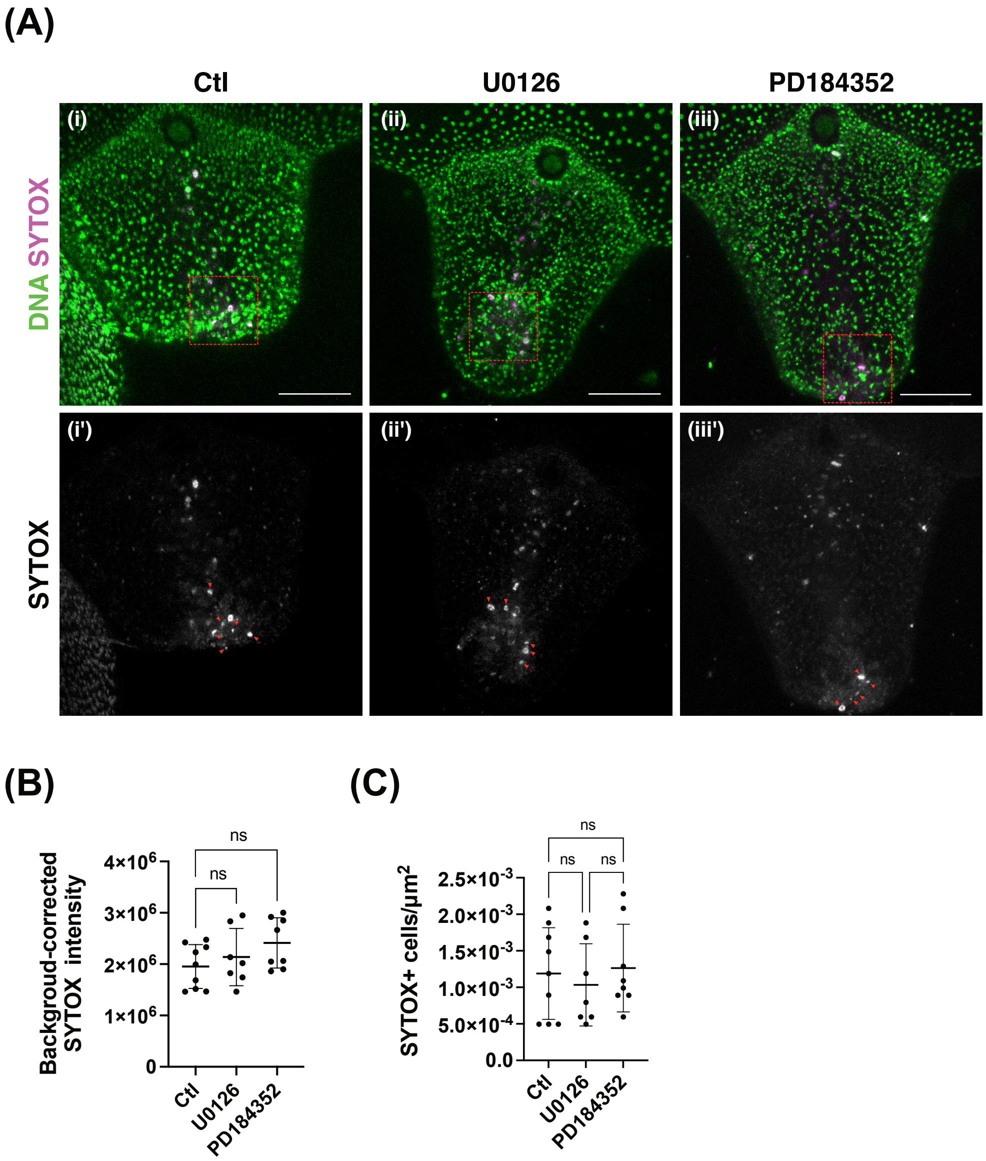

(A)
Ctl
U0126
PD184352
(i)
(ii)
(iii)
DNA SYTOX
(i')
(ii')
(iii')
SYTOX
(B)
Backgroud-corrected SYTOX intensity
4×10^6
3×10^6
2×10^6
1×10^6
0
ns
ns
Ctl
U0126
PD184352
(C)
SYTOX+ cells/µm^2
2.5×10^-3
2.0×10^-3
1.5×10^-3
1.0×10^-3
5.0×10^-4
0.0
ns
ns
ns
Ctl
U0126
PD184352

### Figure S7

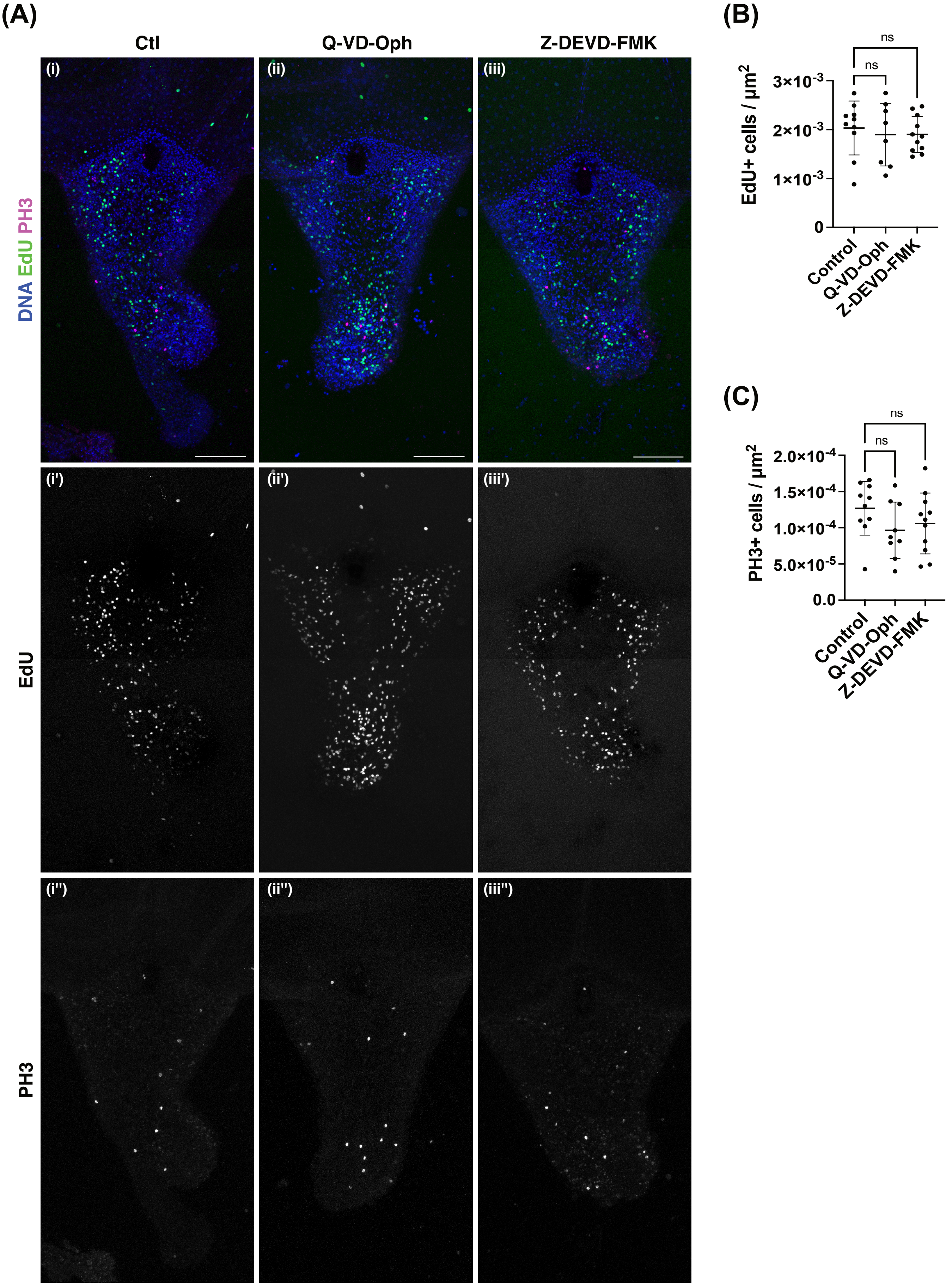
(A)
Ctl
Q-VD-Oph
Z-DEVD-FMK
(i)
(ii)
(iii)
DNA EdU PH3
(i')
(ii')
(iii')
EdU
(i'')
(ii'')
(iii'')
PH3
(B)
EdU+ cells / µm2
3×10-3
2×10-3
1×10-3
0
ns
ns
Control
Q-VD-Oph
Z-DEVD-FMK
(C)
PH3+ cells / µm2
2.0×10-4
1.5×10-4
1.0×10-4
5.0×10-5
0.0
ns
ns
Control
Q-VD-Oph
Z-DEVD-FMK

### Figure S8

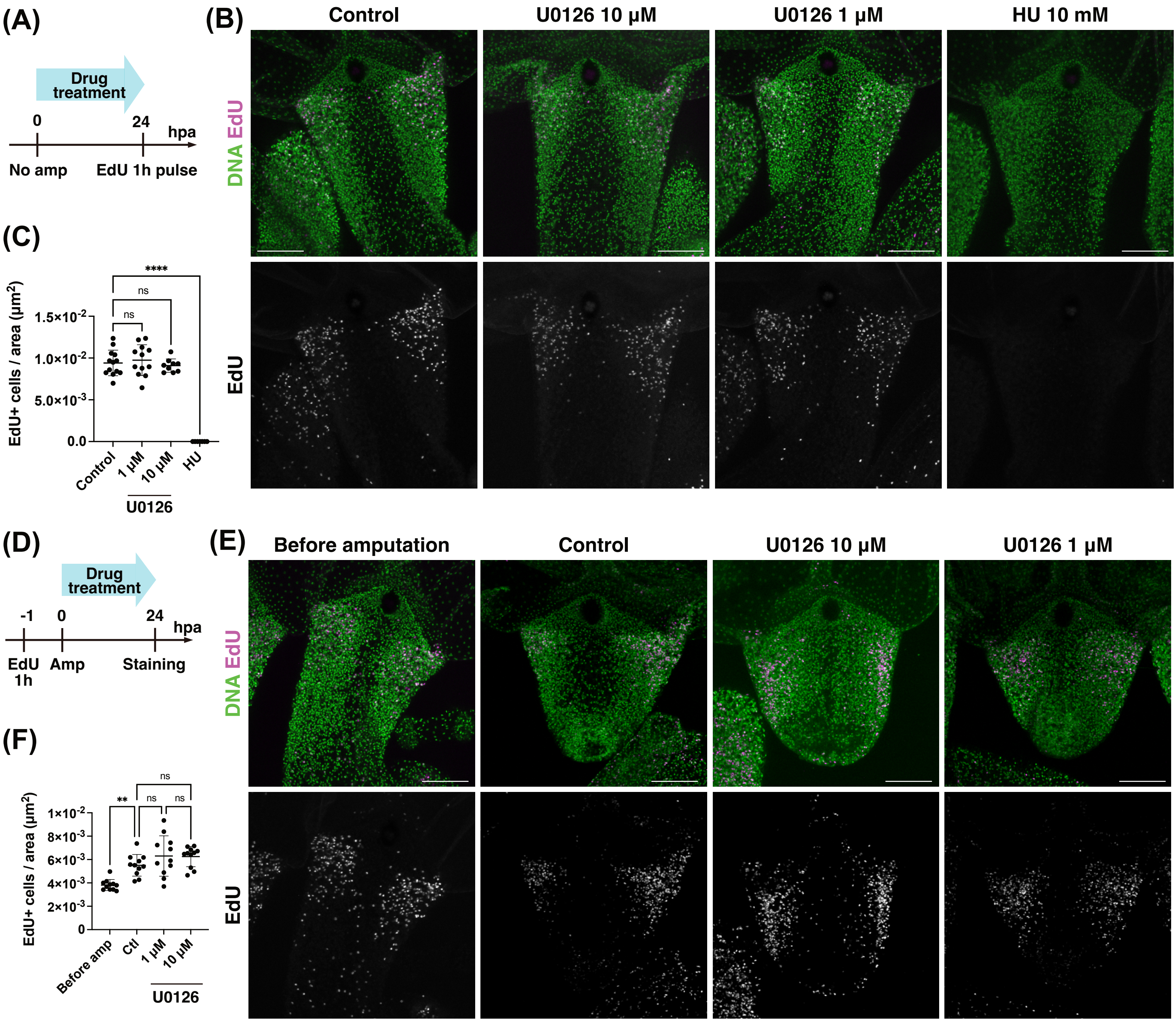

(A)
Drug treatment
0
24
hpa
No amp
EdU 1h pulse
(B)
Control
U0126 10 μM
U0126 1 μM
HU 10 mM
DNA EdU
EdU
(C)
EdU+ cells / area (μm²)
1.5×10⁻²
1.0×10⁻²
5.0×10⁻³
0.0
****
ns
ns
Control
1 μM
10 μM
HU
U0126
(D)
Drug treatment
-1
0
24
hpa
EdU 1h
Amp
Staining
(E)
Before amputation
Control
U0126 10 μM
U0126 1 μM
DNA EdU
EdU
(F)
EdU+ cells / area (μm²)
1×10⁻²
8×10⁻³
6×10⁻³
4×10⁻³
2×10⁻³
0
**
ns
ns
ns
Before amp
Ctl
1 μM
10 μM
U0126
